# TMEM9 and SEC22B interact with ClC-5 to shape renal proximal tubule function and Dent’s Disease type I pathogenesis

**DOI:** 10.1101/2025.11.03.686312

**Authors:** M Durán, A Casal-Pardo, E Sarró, F Stein, S Cases-Palau, C Garcia, G Ariceta, A Meseguer, G Cantero-Recasens

## Abstract

Cl^-^/H^+^ antiporter ClC-5 is a key regulator of renal proximal tubule function, primarily by controlling endosomal acidification. Loss-of-function mutations in ClC-5 cause Dent’s Disease type 1 (DD1), a rare renal tubulopathy that progresses to kidney failure with varying severity. To understand the specific mechanisms linking ClC-5 loss-of-function with DD1 phenotypic heterogeneity, three ClC-5 pathogenic mutations (I524K, E527D and V523Δ) were investigated using renal proximal tubule epithelial cell lines. We have identified distinct intracellular retention patterns of these ClC-5 mutants, all leading to altered endolysosomal pH and disrupted secretory pathway organization. Through an interactome analysis, we have uncovered several ClC-5 partners including TMEM9, which interacted with all ClC-5 forms. TMEM9 knockdown (KD) recapitulated key DD1 characteristics, like defective endocytosis and epithelial dedifferentiation, but paradoxically enhanced endosomal acidification. The secretory pathway was also altered in cells lacking TMEM9 (e.g. enlarged endosomes, and fragmented Golgi apparatus). A similar phenotype was observed in cells depleted of SEC22B; an R-SNARE that we have found to specifically interact with wild-type ClC-5, but not the mutants. Notably, its deletion impaired ClC-5 trafficking, causing its retention at the Golgi and endosomes. Altogether, our findings highlight the relevance of TMEM9 and SEC22B in ClC-5 function, suggesting them as DD1 pathophysiology modifiers.

## Introduction

Dent’s Disease type 1 (DD1) is a rare X-linked renal proximal tubulopathy caused by pathogenic variants in the *CLCN5* gene^1,2^. This disease is characterized by low-molecular-weight proteinuria (LMWP), due to impaired endocytosis of filtered proteins at the proximal tubule. This is often accompanied by hypercalciuria, together with high prevalence of nephrocalcinosis and recurrent nephrolithiasis. Furthermore, DD1 patients may show incomplete Fanconi syndrome, characterized by affected reabsorption of other solutes such as amino acids, glucose or phosphate. Other manifestations, as hypophosphatemic rickets or abnormal electrolyte balance, are detected occasionally. The disease follows a progressive course, leading to chronic kidney disease (CKD) and subsequent renal fibrosis. Renal failure, requiring renal replacement therapy (RRT), occurs in up to 80% of patients, typically by the fourth or fifth decade of life^3–5^. Currently, there is no curative therapy for DD1, and clinical management is supportive, focusing on the reduction of hypercalciuria and the prevention of nephrolithiasis^3^. The notable interpatient variability in clinical phenotype and the age-dependent progression of symptoms present a significant diagnostic challenge in the absence of genetic confirmation to distinguish DD1 from other renal tubulopathies^3,6^.

The *CLCN5* gene encodes the ClC-5 protein, a chloride/proton (Cl^-^/H^+^) antiporter mainly expressed on the endosomal and apical membranes of renal proximal tubule (PT) epithelial cells. ClC-5 function is crucial for the correct acidification of the endolysosomal system, a process essential for receptor-mediated endocytosis and the subsequent recycling of protein receptors and ligands^7^. Loss-of-function mutations in ClC-5 disrupt the endocytic activity, impairing the trafficking of endocyted proteins to lysosomes for degradation and preventing the recycling of key protein receptors. This leads to LMWP, the hallmark of the disease, and contributes to the overall PT dysfunction^7–9^. Furthermore, ClC-5 dysfunction has also been shown to cause epithelial cell dedifferentiation and has been implicated in the development of renal fibrosis^10,11^. Studies in DD1 cell and mouse models have demonstrated that ClC-5 modulates the synthesis and degradation of extracellular matrix components related to the fibrotic tissue, specifically collagen type I and IV^11^. Loss of ClC-5 alters this homeostatic balance, leading to the abnormal accumulation and release of collagens, contributing to the progressive renal fibrosis observed in DD1 patients.

Over 250 different pathogenic variants in *CLCN5* have been identified as the cause of DD1^12^. While a clear genotype-phenotype correlation remains elusive, some associations have been reported. For example, studies have shown that decreased glomerular filtration rate (GFR) is more frequent in DD1 patients with mutations affecting the pore or the CBS domains of ClC-5 compared to those with early-stop mutations^3^. Additionally, protein-truncating variants have been associated with a higher risk of nephrolithiasis and progression to end-stage renal disease (ESRD)^13^. Despite these findings, the specific molecular mechanisms that link *CLCN5* mutations to the broad spectrum of clinical severity and phenotypic heterogeneity in DD1 patients remain to be fully elucidated.

The absence of clear link between ClC-5 mutations and DD1 clinical outcome, and the limited therapeutic options, highlight a significant unmet need for a deeper study of the possible phenotype modifiers. Therefore, we have performed an interactome analysis aiming to identify novel interactors that could explain patient phenotypic variability and serve as potential therapeutic targets for DD1.

## Results

### 1. Pathogenic ClC-5 mutations lead to impaired endolysosomal acidification and deficient intracellular trafficking

Previous work from our group revealed that distinct pathogenic ClC-5 mutations impair endocytosis, which explains LMWP, the hallmark of DD1. However, these mutations were shown to modulate different cellular pathways, leading to variable levels of epithelial dedifferentiation and collagen release^10,11^. This could explain differential disease progression to renal fibrosis and the high phenotypic variability observed among DD1 patients^3^, raising a key question: how do different ClC-5 mutations, all of which disrupt the antiporter function, cause such heterogeneity of cellular responses that may lead to the wide DD1 clinical outcome range?

To investigate the link between ClC-5 mutations and DD1 phenotypic variability, we have used disease cell models previously generated and fully characterized by our group^10^. These cell models are based on the RPTEC/TERT1 renal proximal tubule epithelial cell line, which have been depleted of endogenous *CLCN5* and stably express ClC-5 WT (rWT) or different mutated forms (rV523Δ, rE527D or rI524K). Analyses of ClC-5 mRNA and protein levels confirmed a >90% reduction of *CLCN5* endogenous levels and robust expression of exogenous ClC-5 in differentiated rWT and the three mutant cell lines (**Supplementary Figure 1A and 1B**). Furthermore, study of ClC-5 intracellular localization in these cell lines demonstrated that the WT form can reach the plasma membrane (PM)^10^, after trafficking through endoplasmic reticulum (ER), Golgi apparatus (GA), the trans Golgi network (TGN) and endosomal system; while the mutants are retained in different compartments (**Supplementary Figure 1C**). Our results replicate previous studies from our group and others, showing ER retention of the class I mutant I524K, while the class II mutant E527D can reach the endosomes^10,14^. Interestingly, analysis of the intracellular localization of ClC-5 mutant V523Δ, which remained unclassified with reduced localization at the ER, endosomes and PM compared to WT or other mutants ^10^, showed high colocalization with GA markers (**Figure 1A**). Therefore, our data strongly suggest the existence of a new class of ClC-5 mutations and could provide a reasonable explanation why some DD1 patients have extra clinical outcome resembling Dent’s Disease type 2, which is caused by mutations in the Golgi resident OCRL (Inositol polyphosphate 5-phosphatase)^15–17^. Furthermore, analysis of the secretory pathway revealed that ClC-5 mutants cause fragmentation of the Golgi apparatus (**Figure 1B**), and a reduction on the number and volume of endosomes (**Supplementary Figure 1D**).

**Figure 1.**
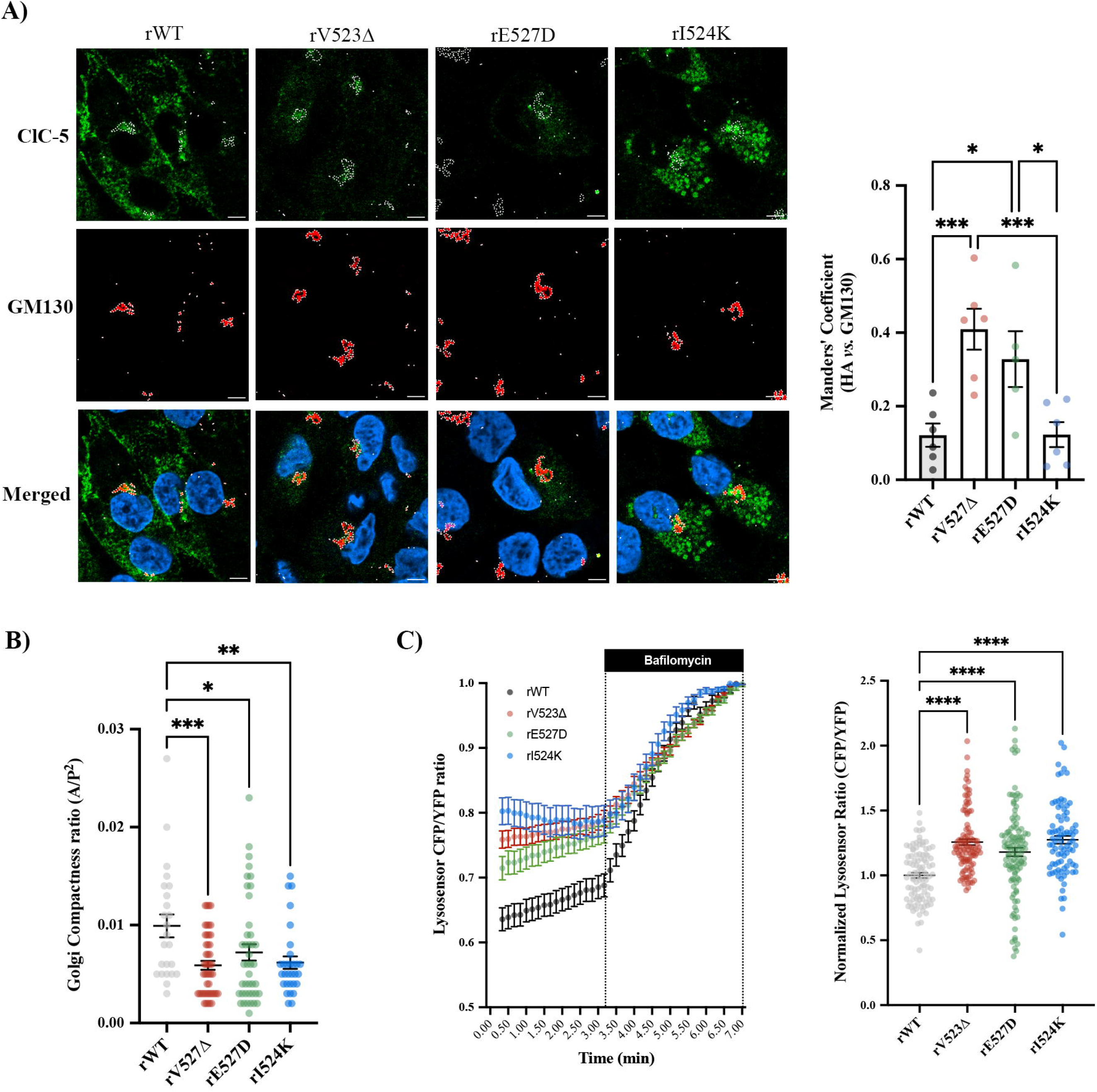
ClC-5 mutants are retained in distinct compartments and lead to secretory pathway alterations. **A)** Representative immunofluorescence z-stack single plane of ClC-5 rWT, rV523Δ, rE527D and rI524K cells stained with anti-HA to detect ClC-5 (green), GM130 (Golgi marker, red) and Hoechst 33342 (blue). Scale bars: 5 µm. Colocalization between ClC-5 and GM130 for all cell lines was calculated from immunofluorescence images by Manders’ coefficient using FIJI (N ≥ 3). Average values ± SEM are plotted as a scatter plot with a bar graph. The y-axis represents Manders’ coefficient of the fraction of ClC-5 (HA) overlapping with GM130. **B)** Quantification of Golgi compactness (ratio between area and perimeter^2^) in rWT, rV523Δ, rE527D and rI524K cells. Data were calculated from immunofluorescence images stained with anti-GM130 using Fiji software. **C)** Time course of the CFP/YFP LysoSensor Yellow/Blue DND-160 dye ratio in ClC-5 rWT, rV523Δ, rE527D and rI524K cells. After 3 min of basal activity, bafilomycin was added as a normalization control. Quantification of the normalized basal CFP/YFP LysoSensor Yellow/Blue DND-160 dye ratio is shown in the right graph. Each dot represents one cell (N ≥ 3). *p<0.05, **p<0.01, ***p<0.001, ****p<0.0001.

Next, considering that ClC-5 is a key regulator of endosomal pH, which impacts endocytic capacity, we used the LysoSensor Yellow/Blue DND-160 dye to analyse the endolysosomal system acidification on our DD1 cell models. Our results showed that, under basal conditions, the three ClC-5 mutant cell lines present a more basic endolysosomal pH than control cells, although E527D expressing cells had a slightly lower pH than the other mutants (pH increase compared to rWT cells, V523Δ: 27%, E527D: 18%, I524K: 27%, p<0.001) (**Figure 1C**). These findings provide an explanation why all these mutants cause defects in endocytosis, as shown previously, but do not account for the differences in the cellular processes and the clinical outcome heterogeneity^3,10,11^.

### 2. Interactome analysis reveal new ClC-5 partners related to the secretory pathway

We have shown that mutations in ClC-5 cause retention of the antiporter in distinct compartments, besides leading to alterations in the secretory pathway, which could explain phenotypic variability in DD1 patients. We postulated that ClC-5 interactors may modulate its function and trafficking, which could be disrupted by pathogenic mutations. To assess this possibility, we performed an interactome analysis of our DD1 cell models expressing the WT or mutated forms of ClC-5 (ClC-5 rWT, rV523Δ, rE527D and rI524K). Our data revealed 210 potential interactors (FDR =< 0.05, logFC => 0.5 compared to control cells), of which 35.7% were specific for ClC-5 rWT (75 proteins), 44.8% were interacting only with mutant forms (49 proteins with rV523Δ, 30 proteins with rE527D, and 30 proteins with rI524K, being 14 of those proteins interacting with two or more mutants), and 4.3% common for all ClC-5 forms (9 proteins) (**Figure 2A**). To uncover the biological significance of ClC-5 interactors, we performed a biological process enrichment analysis using the public database STRING of all potential hits for ClC-5 WT form^18^. Interestingly, five of the top ten processes were related to intracellular protein trafficking, and two were related to the posttranslational modification N-glycosylation, underlying its potential relevance for ClC-5 localization and function (**Figure 2B**). Next, we performed the same enrichment analysis of the lost interactions due to mutations in ClC-5. Interestingly, these WT-specific hits that disappeared for all ClC-5 mutants were related to intracellular protein transport (4 processes) and N-glycosylation (3 processes) (**Figure 2C**). On the other hand, mutations in ClC-5 lead to novel interactions that were grouped in processes associated with protein degradation, phosphorylation or cellular metabolism (**Figure 2D**).

**Figure 2.**
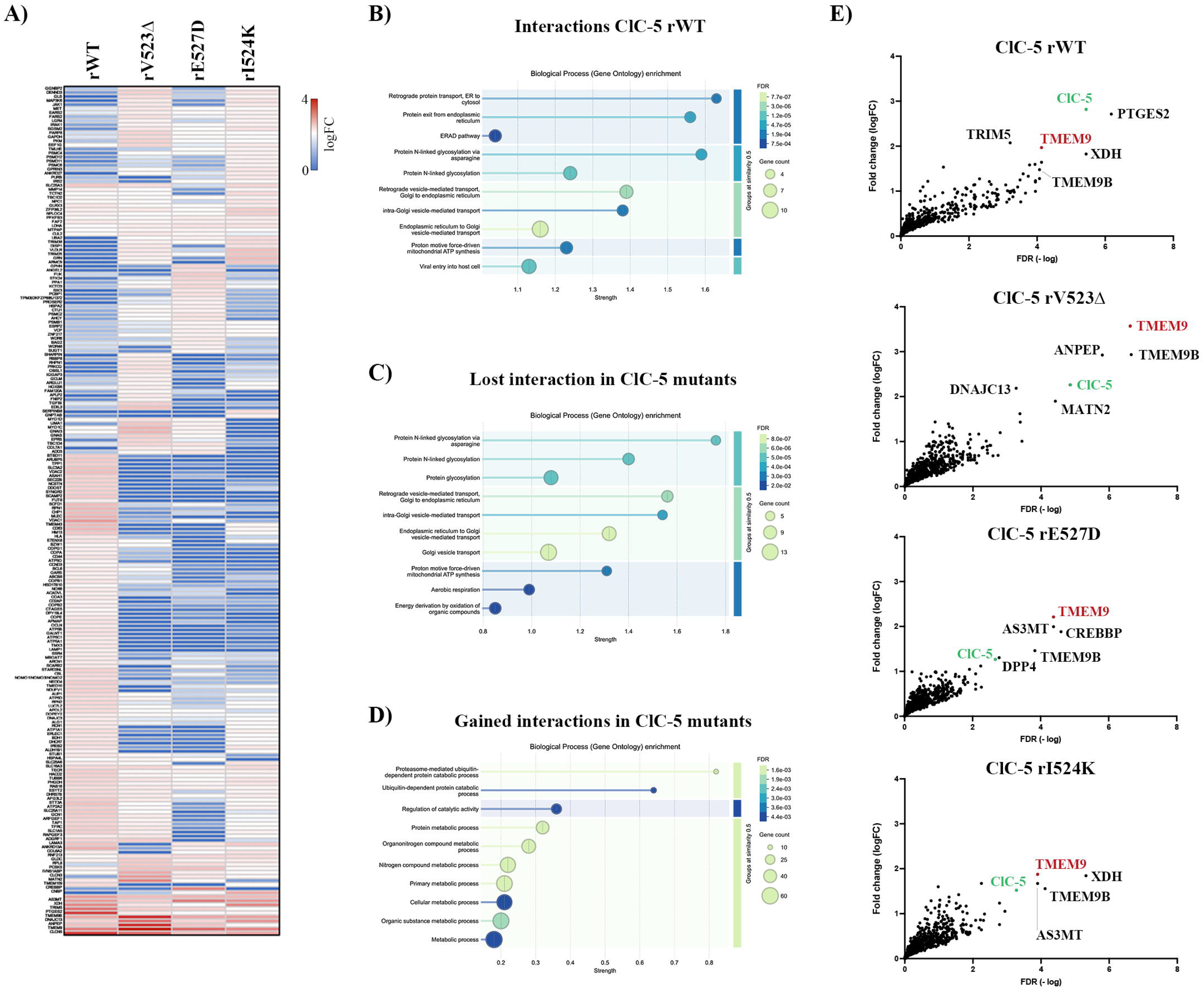
Interactors of ClC-5. **A)** Heatmap showing the levels of the proteins that interact with ClC-5 rWT or distinct mutants with a logFC > |0.5| and an adj.p.val□< □0.05 in comparison to *CLCN5*-silenced cells. **B-D)** Pathway analyses using STRING software of the specific interactors of ClC-5 rWT (**B**), lost interactors in ClC-5 mutants (**C**) or gained interactions by ClC-5 mutants (**D**). **E)** Volcano plot of the interactors identified for each cell line (rWT, rV523Δ, rE527D and rI524K). The y-axis represents the Fold Change of expression levels compared to the CLCN5 silenced cells, and the x-axis represents the FDR (-log). ClC-5 (in green) and top five interactors are named in the plots. TMEM9 is highlighted in red.

Analysis of the interactors for each cell line revealed that all four ClC-5 forms (WT, V523Δ, E527D and I524K) interacted with ClC-5, thus confirming that these mutations do not affect dimerization^19^. Besides, our findings also confirmed TMEM9, the recently identified inhibitory subunit of ClCs^20–22^, as a ClC-5 interactor. Importantly, this interaction was maintained with all ClC-5 mutants, instead of being weakened as previously predicted^22^ (**Figure 2E**). To validate this interaction, we performed co-immunoprecipitation studies in HEK293T cells overexpressing both ClC-5 and TMEM9. As shown in figure 3, our results demonstrated that TMEM9 immunoprecipitated with all forms of ClC-5 (WT, V523Δ, E527D and I524K) (**Figure 3A**). We further studied this interaction in Renal Proximal Tubule Epithelial cells (RPTEC/TERT1) by confocal microscopy. Briefly, rWT, rV523Δ, rE527D and rI524K cells were seeded on glass bottom plates and transfected transiently with TMEM9-Flag after 10-day differentiation. Then, cells were processed for immunofluorescence and imaged by confocal microscopy. Analysis showed strong colocalization of TMEM9 with ClC-5 in all conditions (**Figure 3B**, quantification in **3C**). Notably, we observed a misslocalization of TMEM9 from the lysosomes to other compartments in cells expressing ClC-5 mutants, as shown by a reduction in the colocalization of TMEM9 with the lysosomal marker LAMP1 in ClC-5 mutant cells compared to rWT cells (> 40% reduction, p<0.01) (**Figure 3D**, images in **Supplementary Figure 2A**). Thus, could TMEM9 trafficking impairment due to ClC-5 mutations be leading to some of DD1 characteristics?

**Figure 3.**
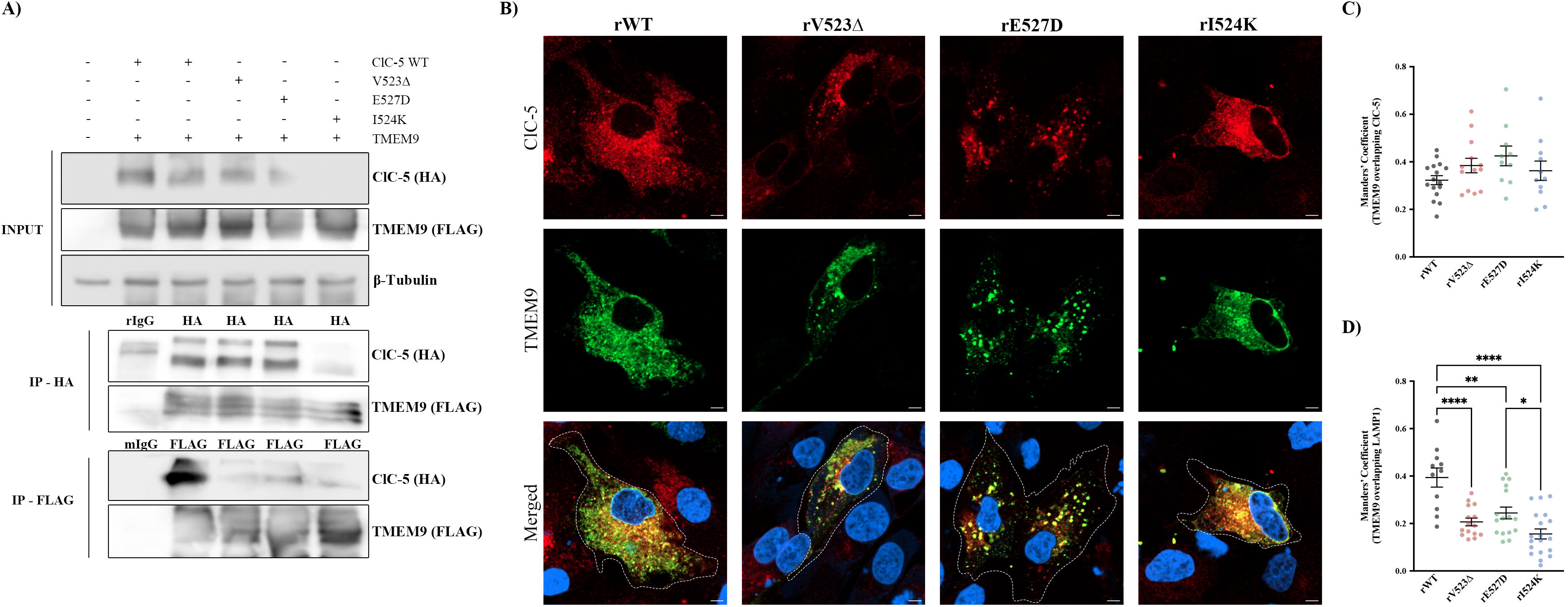
TMEM9 interacts with all forms of ClC-5. **A)** TMEM9 co-immunoprecipitation with the different forms of ClC-5 (WT, rV523Δ, rE527D and rI524K) **B)** Representative immunofluorescence z-projection images of rWT, rV523Δ, rE527D and rI524K cells stained with anti-FLAG (TMEM9) and anti-HA (ClC-5) antibodies. Scale bars: 5 µm. **C)** Colocalization between ClC-5 and TMEM9 for all cell lines was calculated from immunofluorescence images by Manders’ coefficient using FIJI (N ≥ 3). Average values ± SEM are plotted as a scatter plot. The y-axis represents Manders’ coefficient of the fraction of TMEM9 overlapping with ClC-5. **D)** Colocalization between LAMP1 and TMEM9 for all cell lines was calculated from immunofluorescence images by Manders’ coefficient using FIJI (N ≥ 3). Average values ± SEM are plotted as a scatter plot. The y-axis represents Manders’ coefficient of the fraction of TMEM9 overlapping with LAMP1. *p<0.05, **p<0.01, ***p<0.001, ****p<0.0001.

### 3. TMEM9 depletion phenocopies ClC-5 loss-of-function in RPTEC cells

To understand the contribution of TMEM9 to DD1 phenotype, we generated an RPTEC cell line stably depleted of TMEM9 (TMEM9 KD). To confirm knockdown efficiency, RNA was extracted from differentiated TMEM9 KD cells and *TMEM9* expression measured by RT-qPCR. Compared to control cells, TMEM9 KD cells showed a 85% reduction in *TMEM9* mRNA levels (**Supplementary Figure 2B**). Then, we analysed epithelial differentiation by studying the levels of E-cadherin and mucin-1 (MUC1) by WB in cell lysates extracted from control and TMEM9 KD cells. Our results showed a strong reduction of both markers in TMEM9 KD cells compared to control cells (**Figure 4A** and **4B**, respectively). Confocal microscopy study also confirmed MUC1 reduction in TMEM9 KD cells (**Figure 4C**). We next explored whether this reduction in epithelial markers correlated with other dedifferentiation characteristics as seen in cells lacking a functional ClC-5 ^10,11^, such as substrate adhesion, endocytic capacity, or collagen production and release.

**Figure 4.**
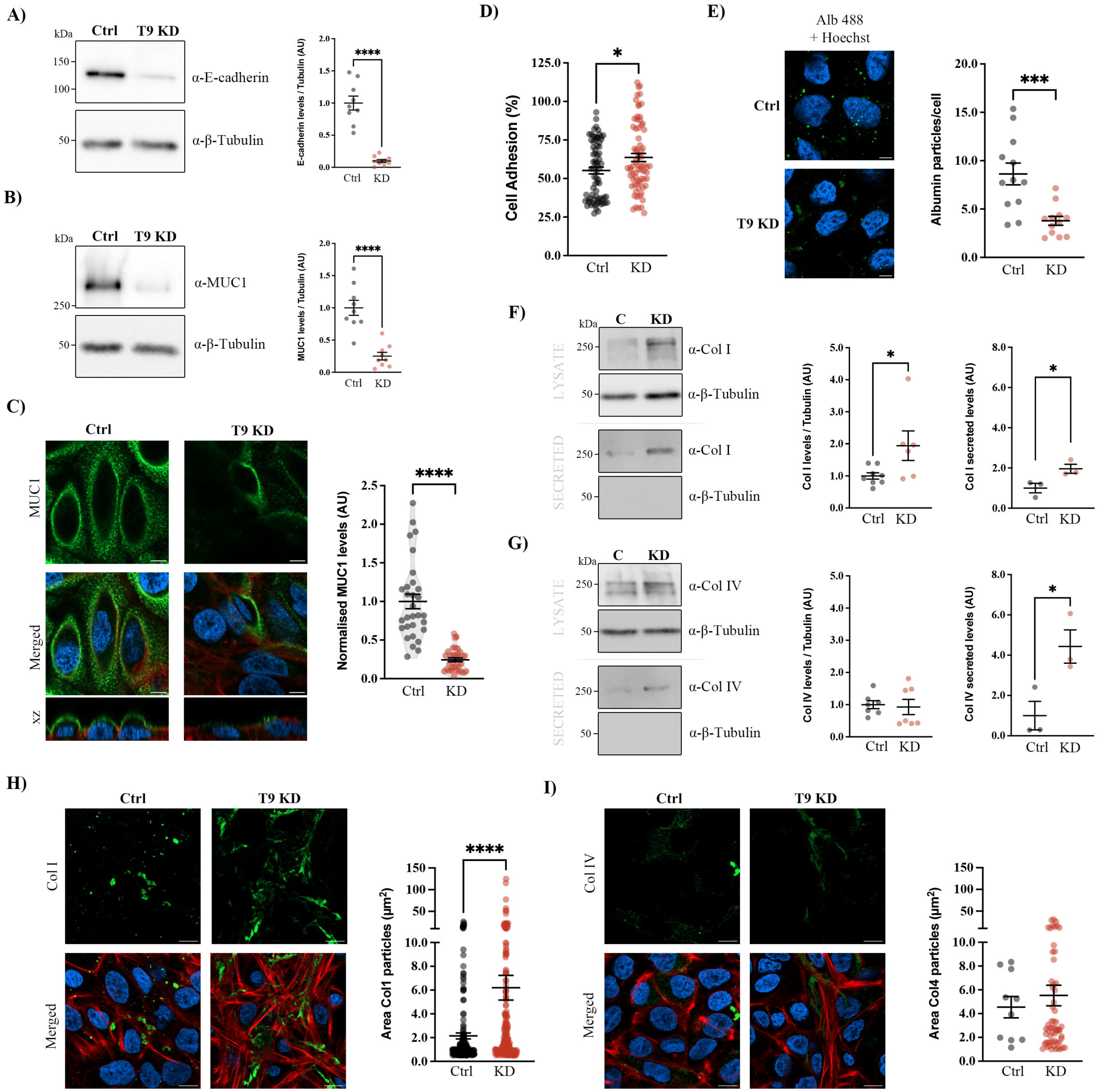
Depletion of TMEM9 leads to proximal tubule epithelial cell dedifferentiation. **A)** E-cadherin protein levels were analysed by Western blot in cell lysates from control or TMEM9 KD cells. Tubulin was used as a loading control. Average values ± SEM are plotted as a scatter plot (N ≥ 3). **B)** Mucin-1 (MUC1) protein levels were analysed by Western blot in cell lysates from control or TMEM9 KD cells. Tubulin was used as a loading control. Average values ± SEM are plotted as a scatter plot (N ≥ 3). **C)** Immunofluorescence z-stack single plane images of control or TMEM9 KD cells stained with anti-MUC1 antibody (green), phalloidin (red), and Hoechst 33342 (blue). Scale bars: 5 μm. **D)** Cell-to-substrate adhesion as assessed by the ability of control or TMEM9 KD cells to bind to tissue culture substrate. **E)** Representative immunofluorescence z-stack single plane images of control and TMEM9 KD cells treated with Alexa Fluor 488-conjugated Albumin (green) stained with Hoechst 33342 (blue). Quantification of albumin uptake was performed by measuring the number of albumin particles/cell in control or TMEM9 KD cells. (N □≥ □3). **F-G)** Collagen type I (Col I) **(F)** or collagen type IV (Col IV) **(G)** protein levels in the lysates or secreted media of control or TMEM9 KD cells. Tubulin was used as a loading control in the lysates, and as a control of lysate contamination in secreted media. The right graph shows the quantification of Col I and Col IV intracellular and secreted levels (N ≥ 3). **H-I)** Representative immunofluorescence z-projection of control or TMEM9 KD cells stained with anti-Col I **(H)** or anti-Col IV **(I)** antibody (green), phalloidin (red), and Hoechst 33342 (blue). Scale bars: 5 μm. Abbreviations: Ctrl: Control cells, T9 KD: TMEM9 KD cells, KD: TMEM9 KD cells, MUC1: Mucin-1, Col I: Collagen type I, Col IV: Collagen type IV, Alb 488: Albumin 488. *p<0.05, **p<0.01, ***p<0.001, ****p<0.0001.

First, our findings revealed enhanced cell-to-substrate adhesion of TMEM9 KD cells compared to control cells (p<0.05) (**Figure 4D**). Second, study of endocytosis using Alexa Fluor 488 labelled albumin showed that TMEM9 deletion strongly reduced albumin uptake compared to control cells (particles/cell: control cells = 8.63, TMEM9 KD = 3.79, p<0.001) (**Figure 4E**). Third, we analysed whether intracellular and extracellular levels of collagen type I and IV (Col I and Col IV), which are key components of renal fibrotic tissues, were affected by TMEM9 depletion, as observed in cells depleted of ClC-5^11^. Our results showed that TMEM9 deletion causes an increase in both intracellular and extracellular Col I levels compared to control cells (2-fold increase, p<0.05), while only Col IV release to the extracellular medium, but not its intracellular expression, was enhanced (4.5-fold increase, p<0.05) (**Figure 4F** and **4G**, respectively). To confirm this specific increase of Col I intracellular levels but not Col IV, we imaged control and TMEM9 KD cells by confocal microscopy. Our data showed that TMEM9 KD cells presented an increase in intracellular Col I levels, which was not observed in cells stained against Col IV (**Figure 4H** and **4I**, respectively). Altogether, our findings showed that lack of TMEM9 promotes epithelial dedifferentiation in RPTEC, similar to the ClC-5 loss-of-function phenotype.

Next, we assessed whether TMEM9 was modulating epithelial dedifferentiation and collagen production through the β-catenin pathway as previously shown for ClC-5^11^. Analysis of β-catenin localization cells showed small increase in its nuclear translocation in TMEM9 KD cells compared to control cells (4% vs 9%, p<0.01) (**Supplementary Figure 3A**). In addition, we studied the transcription of two well-known β-catenin target genes (*BMP4* and *EFNB1*). Our data showed that *BMP4* mRNA levels were up-regulated while *EFNB1* expression was decreased in TMEM9 KD cells (**Supplementary Figure 3B**). Next, we assessed whether Col I and Col IV mRNA levels, which are partially controlled by β-catenin pathway, were altered in TMEM9 KD cells, which could explain the increase in their protein levels (as shown in **Figure 4**). Interestingly, Col I mRNA levels (*COL1A1*) were slightly upregulated in cells depleted of TMEM9 (1.9-fold increase, p<0.05), while Col IV expression was not affected (**Supplementary Figure 3C** and **3D**). These data suggest that other pathways besides β-catenin differentially regulate Col I and Col IV intracellular levels, which could be affected by TMEM9, such as lysosomal degradation or protein trafficking.

### 4. Deletion of TMEM9 affects secretory pathway organization in RPTEC cells

ClC-5 is a key element for maintaining correct endolysosomal pH by positively regulating the v-ATPase, and TMEM9 has recently emerged as a negative regulatory subunit of several ClC antiporters^20,22^. Therefore, we studied the role of TMEM9 in endolysosomal acidification, which could explain some of the dedifferentiation phenotype and its contribution to DD1-like phenotype. Our results showed that TMEM9 KD cells have a lower endolysosomal pH than control cells (20% reduction, p<0.0001) (**Figure 5A**), confirming the inhibitory role of TMEM9 on ClC-5. This could explain why there is not an intracellular accumulation of collagen as observed in cells lacking ClC-5, since lower lysosomal pH could lead to enhanced degradation of collagens. However, it cannot account for the other characteristics which resemble ClC-5 loss-of-function (which lead to higher endolysosomal pH). Therefore, we postulated that lack of TMEM9, by altering endosomal acidification, could affect the organization of the secretory pathway, therefore explaining the defects on endocytosis, protein recycling, and also on ß-catenin pathway activation.

**Figure 5.**
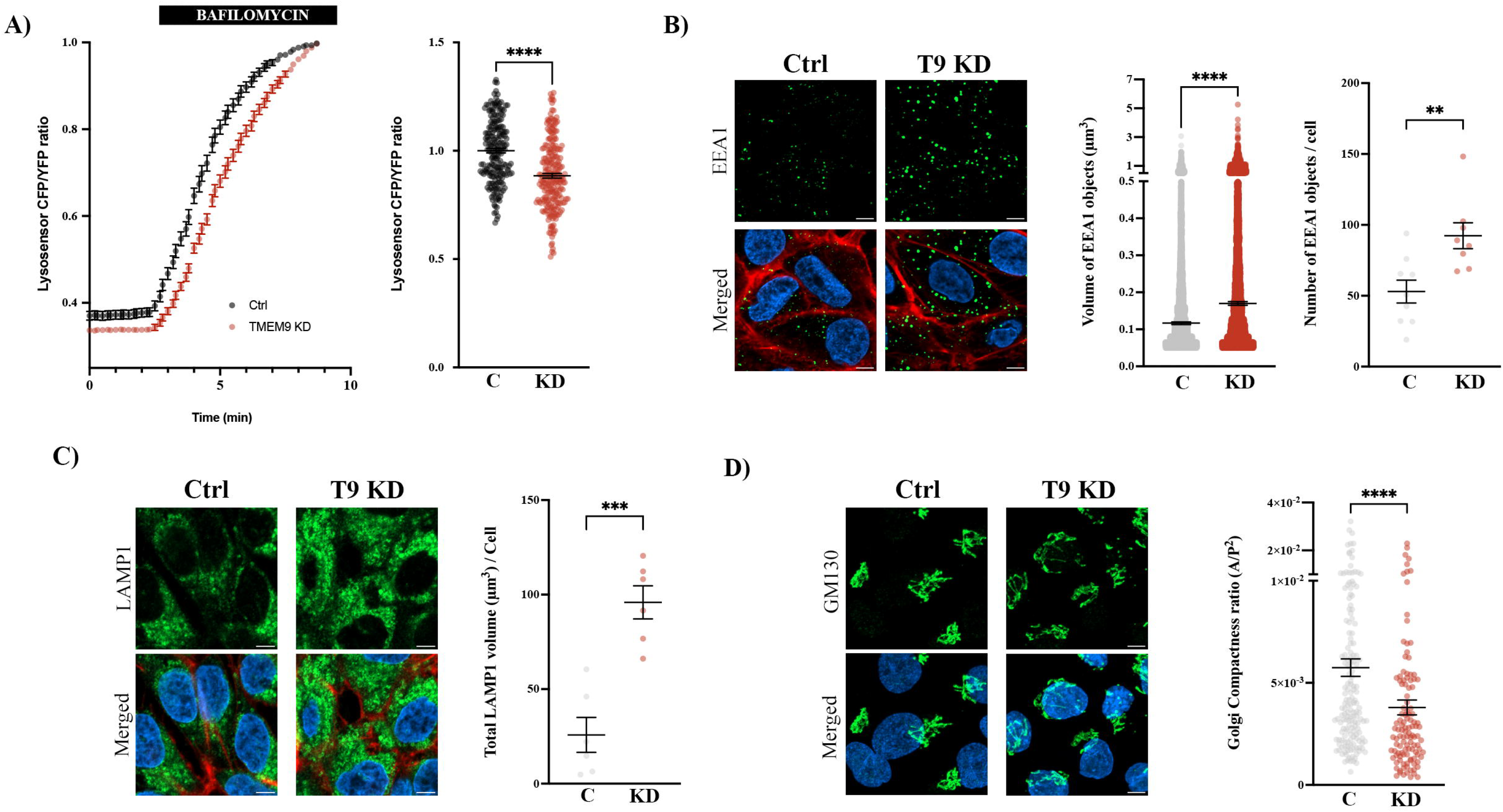
Effect of TMEM9 KD on the secretory pathway. **A)** Time course of the CFP/YFP LysoSensor Yellow/Blue DND-160 dye ratio in ClC-5 control and TMEM9 KD cells. After 3 min of basal activity, bafilomycin was added as a normalization control. Quantification of the normalized basal CFP/YFP LysoSensor Yellow/Blue DND-160 dye ratio is shown in the left graph. Each dot represents one cell (N ≥ 3). **B)** Immunofluorescence z-stack single planes of control or TMEM9 KD cells stained with anti-EEA1 (endosomal marker, green), phalloidin (red) and Hoechst 33342 (blue). Scale bars: 5 µm. Volume and number of endosomes were quantified from immunofluorescence images using Fiji software (N ≥ 3). **C)** Immunofluorescence z-stack single planes of control or TMEM9 KD cells stained with anti-LAMP1 (lysosomal marker, green), phalloidin (red) and Hoechst 33342 (blue). Scale bars: 5 µm. Lysosomal volume was quantified from immunofluorescence images using Fiji software (N ≥ 3). **D)** Representative z-projection images of control and TMEM9 KD cells stained with anti-GM130 antibody (green). Quantification of Golgi compactness (ratio between area and perimeter^2^) in control and TMEM9 KD cells. Data were calculated from immunofluorescence images stained with anti-GM130 using Fiji software (N ≥ 3). Abbreviations: Ctrl: Control cells, C: Control cells, T9 KD: TMEM9 KD cells, KD: TMEM9 KD cells. **p<0.01, ***p<0.001, ****p<0.0001.

To assess this, we studied the secretory pathway by confocal microscopy using markers of the endosomes (EEA1), lysosomes (LAMP1), and Golgi apparatus (GM130). Immunofluorescence analysis of EEA1 revealed that TMEM9 KD cells presented more endosomes (control: 50 EEA1 objects/cell, TMEM9 KD: 80 EEA1 objects/cell, p<0.01), which were larger compared to those of control cells (0.12 µm^3^ vs 0.17 µm^3^, respectively, p<0.0001) (**Figure 5B**). Similarly, we found increased levels of LAMP1 covered volume in TMEM9 depleted cells compared to control cells (90 µm^3^ vs 30 µm^3^, p<0.001) (**Figure 5C**). Surprisingly, analysis of GM130 showed a reduced compactness of the Golgi apparatus in TMEM9 KD cells (**Figure 5D**) that resembles the effect observed in cells expressing ClC-5 mutants (**Figure 1B**). Thus, how are TMEM9 or ClC-5 loss-of-function mutations causing a disruption of the secretory pathway organization and specially the Golgi apparatus? Deep literature review of the interactome hits related to intracellular trafficking showed that an R-SNARE called SEC22B, which only interacts with ClC-5 WT form, can affect GA organization and its loss-of-function leads to GA fragmentation^23^. Interestingly, one study on human soluble protein complexes identified SEC22B as a potential interactor of TMEM9^24^. Therefore, it is possible that SEC22B interaction with TMEM9 and ClC-5 could be necessary for their correct trafficking and function.

### 5. SEC22B interacts with ClC-5 WT, but not with ClC-5 mutants

Our analysis showed that SEC22B specifically interacts with ClC-5 WT, while this interaction is lost in the mutants, which could explain some of their trafficking defects. To confirm this interaction, we performed a co-immunoprecipitation experiment co-transfecting SEC22B and the different forms of ClC-5 (WT, V523Δ, E527D and I524K) in HEK293T cells. Our results confirmed that only ClC-5-WT is capable of interacting with SEC22B (**Supplementary Figure 4A**). However, it is possible that ClC-5 mutations cause a decrease in SEC22B levels, which may explain the loss of this interaction. To exclude this, mRNA and protein was extracted from differentiated RPTEC ClC-5 rWT, rV523Δ, rE527D and rI5234K cells and SEC22B expression analysed. Importantly, our data showed no major differences between WT or mutant cell lines regarding SEC22B mRNA or protein levels (**Figure 6A** and **6B**, respectively). Next, we further studied SEC22B and ClC-5 interaction in RPTEC cells by confocal microscopy. We transiently transfected our DD1 cell lines (rWT, rV523Δ, rE527D or rI524K) with SEC22B. As shown in Figure 6D, SEC22B nicely colocalized with ClC-5 WT form, but not with any of the ClC-5 mutants (**Figure 6C**, quantification in **6D**). Next, we studied SEC22B intracellular localization in the different cell lines, in order to understand whether expression of ClC-5 mutants could lead to trafficking defects of SEC22B. Colocalization analyses with KDEL (ER marker) revealed that SEC22B localization was not altered by the expression of ClC-5 mutants compared to ClC-5 WT (**Figure 6E**, images in **Supplementary Figure 4B**).

**Figure 6.**
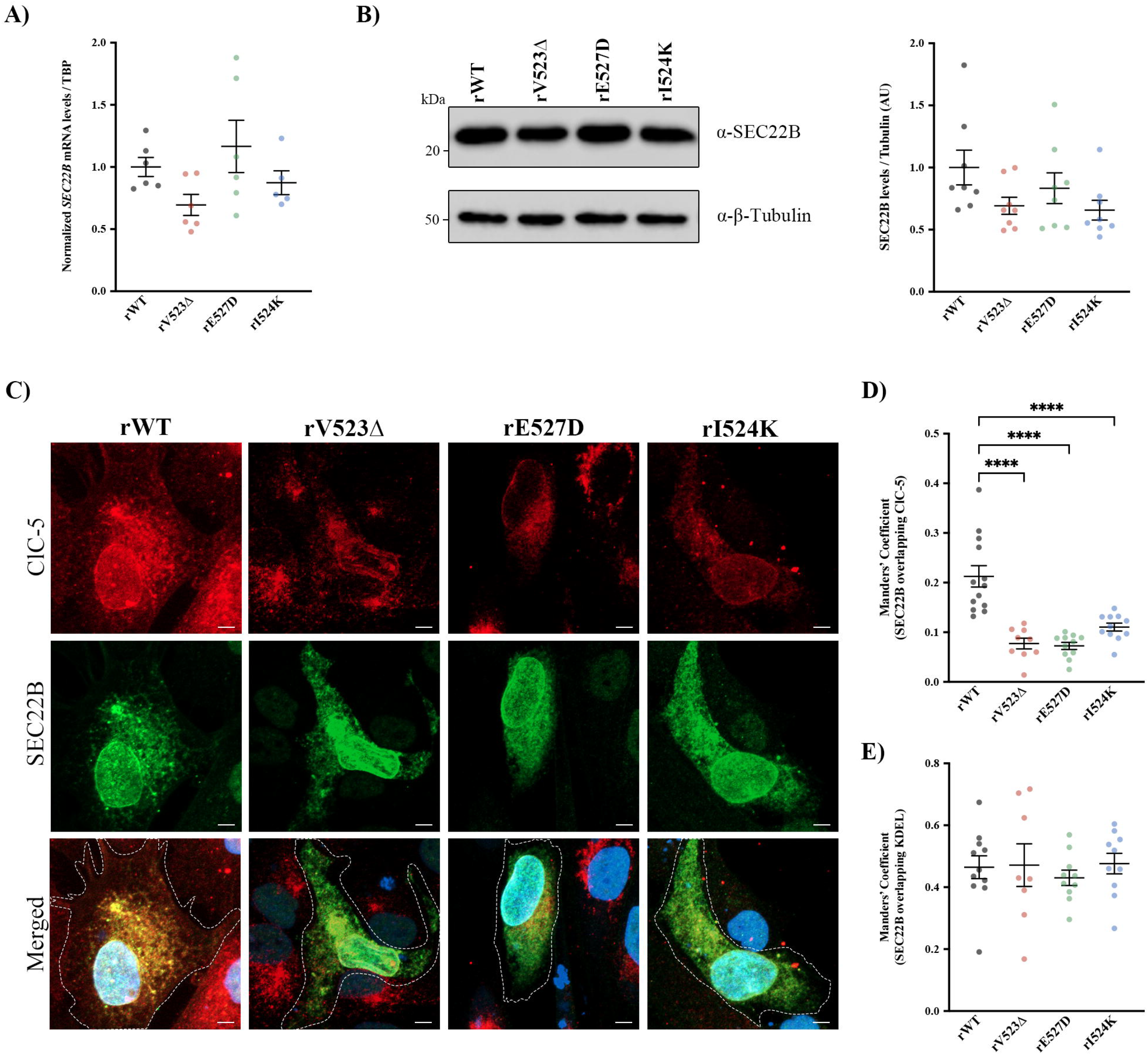
SEC22B specifically interacts with ClC-5 WT. **A)** Endogenous *SEC22B* mRNA levels normalized to values of *TBP* from in rWT, rV523Δ, rE527D and rI524K cells. All values are represented as a relative value compared with rWT cells. Average values ± SEM are plotted as a scatter plot with a bar graph (N ≥ 3). **B)** Cell lysates from in rWT, rV523Δ, rE527D and rI524K cells were analysed by Western blot with an anti-SEC22B to test expression levels. Tubulin was used as a loading control. Average values ± SEM are plotted as a scatter plot (N ≥ 3). **C)** Representative z-projection images of rWT, rV523Δ, rE527D and rI524K cells stained with anti-Flag (SEC22B) and anti-HA (ClC-5) antibody. Scale bars: 5 µm. **D)** Colocalization between ClC-5 and SEC22B for all cell lines was calculated from immunofluorescence images by Manders’ coefficient using FIJI (N ≥ 3). Average values ± SEM are plotted as a scatter plot. The y-axis represents Manders’ coefficient of the fraction of SEC22B overlapping with ClC-5 (HA). **E)** Colocalization between SEC22B and KDEL (ER marker) for all cell lines was calculated from immunofluorescence images by Manders’ coefficient using FIJI (N ≥ 3). Average values ± SEM are plotted as a scatter plot. The y-axis represents Manders’ coefficient of the fraction of SEC22B overlapping with KDEL signal. ****p<0.0001.

### 6. SEC22B modulates RPTEC secretory pathway

SEC22B is an R-SNARE that regulates anterograde and retrograde trafficking ER-Golgi. Notably, deletion of SEC22B has shown to alter the transport of proteins from the Golgi to the PM, possibly due to Golgi fragmentation^23^. Therefore, could SEC22B have a role in intracellular protein trafficking of proximal tubule epithelial cells, controlling ClC-5 transport which may then impact proximal tubule function and explain DD1 phenotypic variability? To answer this question, we have generated an RTPEC cell line stably depleted of SEC22B (SEC22B KD). mRNA and protein were extracted from control and SEC22B KD cells to evaluate the knockdown efficiency. Our results confirmed a higher than 85% reduction of SEC22B at both RNA and protein levels (**Supplementary Figure 4C** and **4D**, respectively). Next, we characterized the effect of SEC22B deletion in the secretory pathway by imaging cells with confocal microscopy. Our data confirmed a major fragmentation of the GA (**Figure 7A**), an increase in the volume and number of endosomes (**Figure 7B**), and higher LAMP1 total volume (**Figure 7C**) in SEC22B KD cells compared to control cells. Then, we studied distinct parameters related to proximal tubule dysfunction observed in DD1, such as endocytosis, epithelial differentiation or collagen production. Importantly, endocytosis of albumin labelled particles was strongly reduced in SEC22B KD cells compared to control cells (4 vs. 9 particles/cell, respectively, p<0.0001) (**Figure 7D**). Moreover, expression levels of E-cadherin and MUC1, two epithelial markers, were decreased in cells depleted of SEC22B (**Figure 7E and 7F**, respectively), suggesting epithelial dedifferentiation similar to our results in ClC-5 mutant or TMEM9 KD cells. Surprisingly, intracellular levels of collagen I or IV were not affected, and only secretion of collagen I was higher in SEC22B KD cells than control cells (1.8 fold increase, p<0.05) (**Figure 7G-H**). Accordingly, Col I (*COL1A1*) and IV (*COL4A1*) mRNA levels were not affected by SEC22B deletion (**Supplementary Figure 5A-B**), and our results revealed that β-catenin pathway was not altered in SEC22B KD cells, as shown by the lack of enhanced nuclear translocation (**Supplementary Figure 5C**) or affected transcription of other target genes (**Supplementary Figure 5D**).

**Figure 7.**
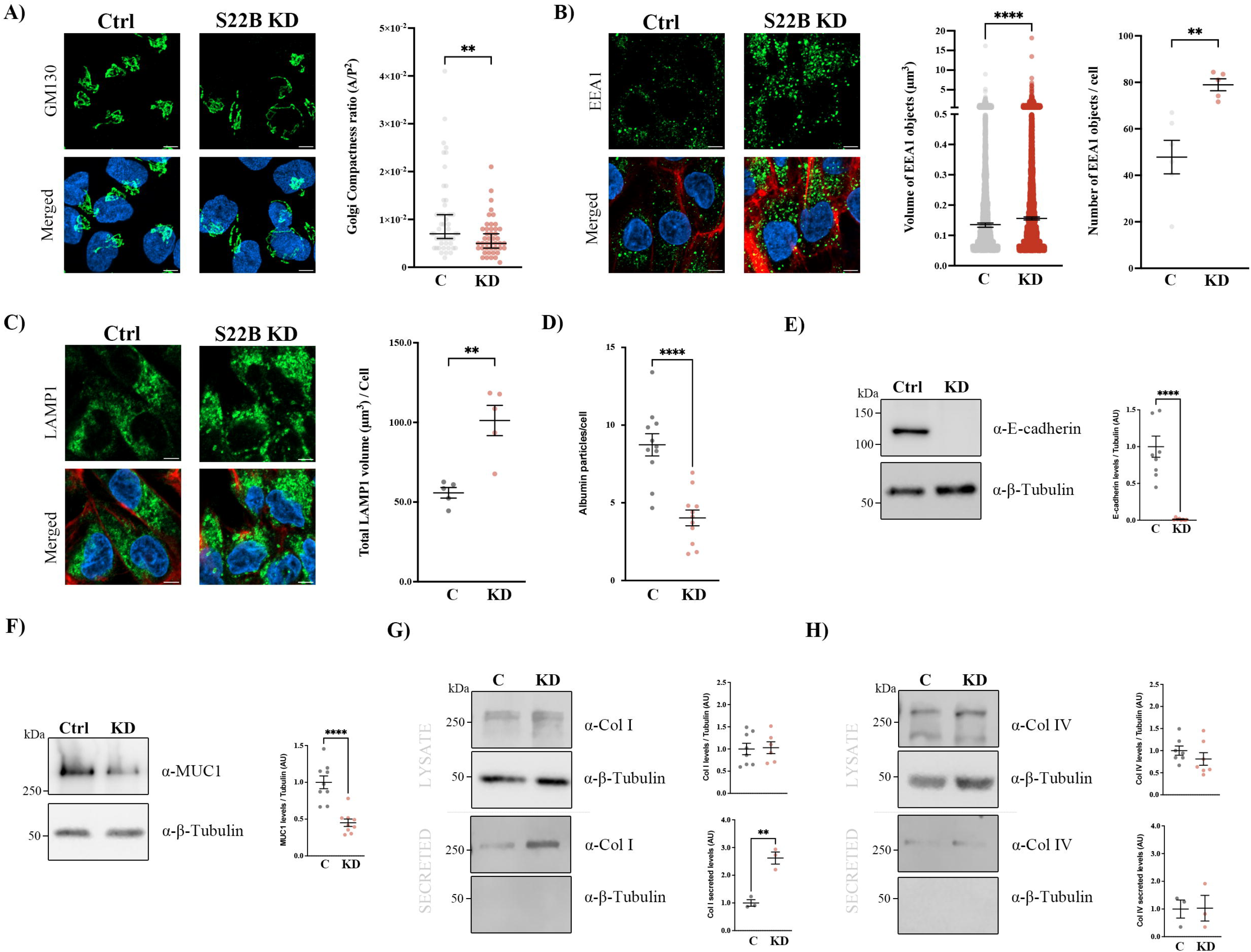
SEC22B depletion disrupts the secretory pathway of RPTEC cells. **A)** Representative z-projection images of control and SEC22B KD cells stained with anti-GM130 antibody. Quantification of Golgi compactness (ratio between area and perimeter^2^) in control and SEC22B KD cells. Data were calculated from immunofluorescence images stained with anti-GM130 using Fiji software. **B)** Immunofluorescence z-stack single planes of control or SEC22B KD cells stained with anti-EEA1 (endosomal marker, green), phalloidin (red) and Hoechst 33342 (blue). Scale bars: 5 µm. Volume and number of endosomes were quantified from immunofluorescence images using Fiji software (N ≥ 3). **C)** Immunofluorescence z-stack single planes of control or SEC22B KD cells stained with anti-LAMP1 (lysosomal marker, green), phalloidin (red) and Hoechst 33342 (blue). Scale bars: 5 µm. Lysosomal volume was quantified from immunofluorescence images using Fiji software (N ≥ 3). **D)** Quantification of albumin uptake was performed by measuring the number of albumin particles/cell in control or SEC22B KD cells (N □≥ 3). **E)** E-cadherin protein levels were analysed by Western blot in cell lysates from control or SEC22B KD cells. Tubulin was used as a loading control. Average values ± SEM are plotted as a scatter plot (N ≥ 3). **F)** Mucin-1 (MUC1) protein levels were analysed by Western blot in cell lysates from control or SEC22B KD cells. Tubulin was used as a loading control. Average values ± SEM are plotted as a scatter plot (N ≥ 3). **G-H)** Collagen type I (Col I) (**G**) or collagen type IV (Col IV) (**H**) protein levels in the lysates or secreted media of control or SEC22B KD cells. Tubulin was used as a loading control in the lysates, and as a control of lysate contamination in secreted media. The right graph shows the quantification of Col I and Col IV intracellular and secreted levels (N ≥ 3). Abbreviations: Ctrl: Control cells, C: Control cells, S22B KD: SEC22B KD cells, KD: SEC22B KD cells. *p<0.05, **p<0.01, ***p<0.001, ****p<0.0001.

### 7. SEC22B controls intracellular trafficking of ClC-5

Considering that 1) SEC22B interacts only with ClC-5 WT, 2) SEC22B is a SNARE involved in intracellular trafficking, and 3) ClC-5 mutants are retained in different compartments, we postulated that SEC22B was necessary for ClC-5 correct targeting to its final destination. Therefore, the loss of SEC22B could impair this process, which could explain the observed decrease in endocytosis and epithelial markers (hallmarks of ClC-5 loss-of-function and DD1).

To investigate this possibility, and understand the differences between SEC22B and TMEM9 relationship with ClC-5, we analysed the localization of ClC-5 by confocal microscopy. Differentiated cells lacking SEC22B or TMEM9 and their respective controls were transiently transfected with ClC-5 WT, processed for immunofluorescence analysis and co-stained with phalloidin (PM), anti-GM130 (GA), anti-EEA1 (endosomes) or anti-KDEL (ER). Our results showed that depletion of SEC22B caused a retention of ClC-5 at the GA and endosomes, as revealed by lower colocalization with PM marker and higher colocalization with GM130 and EEA1 (**Figure 8A**). Importantly, exit from ER was not affected by SEC22B deletion (ClC-5 vs KDEL colocalization, control: 0.13, SEC22B KD: 0.15, n.s.). On the other hand, our data clearly showed that TMEM9 KD did not affect intracellular localization of ClC-5, being it able to normally traffic to the PM in both control and TMEM9 KD cells (**Figure 8B**). Then, we studied whether endolysosomal acidification was affected in SEC22B KD cells compared to control cells. Surprisingly, endolysosomal pH was lower in cells lacking SEC22B (**Figure 8C**), similar to the effect observed in TMEM9 KD cells. We suggest this could be a consequence of the retention of ClC-5 in endosomes, which would promote higher activity of vATPase, therefore leading to lower pH. Furthermore, we analysed whether SEC22B depletion affected TMEM9 trafficking. We overexpressed TMEM9 in control and SEC22B KD cells, and analysed its intracellular localization. Notably, our findings revealed a reduction in TMEM9 localization at the lysosomes, as shown by a decreased colocalization with the lysosomal marker LAMP1 (Manders’ coefficient: 0.33 vs 0.21, p<0.05) (**Supplementary Figure 6A**). This, together with the increased localization of ClC-5 at endosomes, may explain the higher acidification observed in SEC22B KD cells. Finally, we analysed ClC-5 endogenous levels in SEC22B KD, TMEM9 KD and respective control cells. Our results showed no effect on ClC-5 mRNA levels in SEC22B KD cells, and a small reduction in TMEM9 KD cells compared to respective control cells (**Supplementary Figure 6B)**. Accordingly, analysis of ClC-5 protein levels showed only a reduction of ClC-5 levels in TMEM9 KD cells, which we suggest could be a consequence of epithelial cell dedifferentiation and trafficking defects (**Supplementary Figure 6C**).

**Figure 8.**
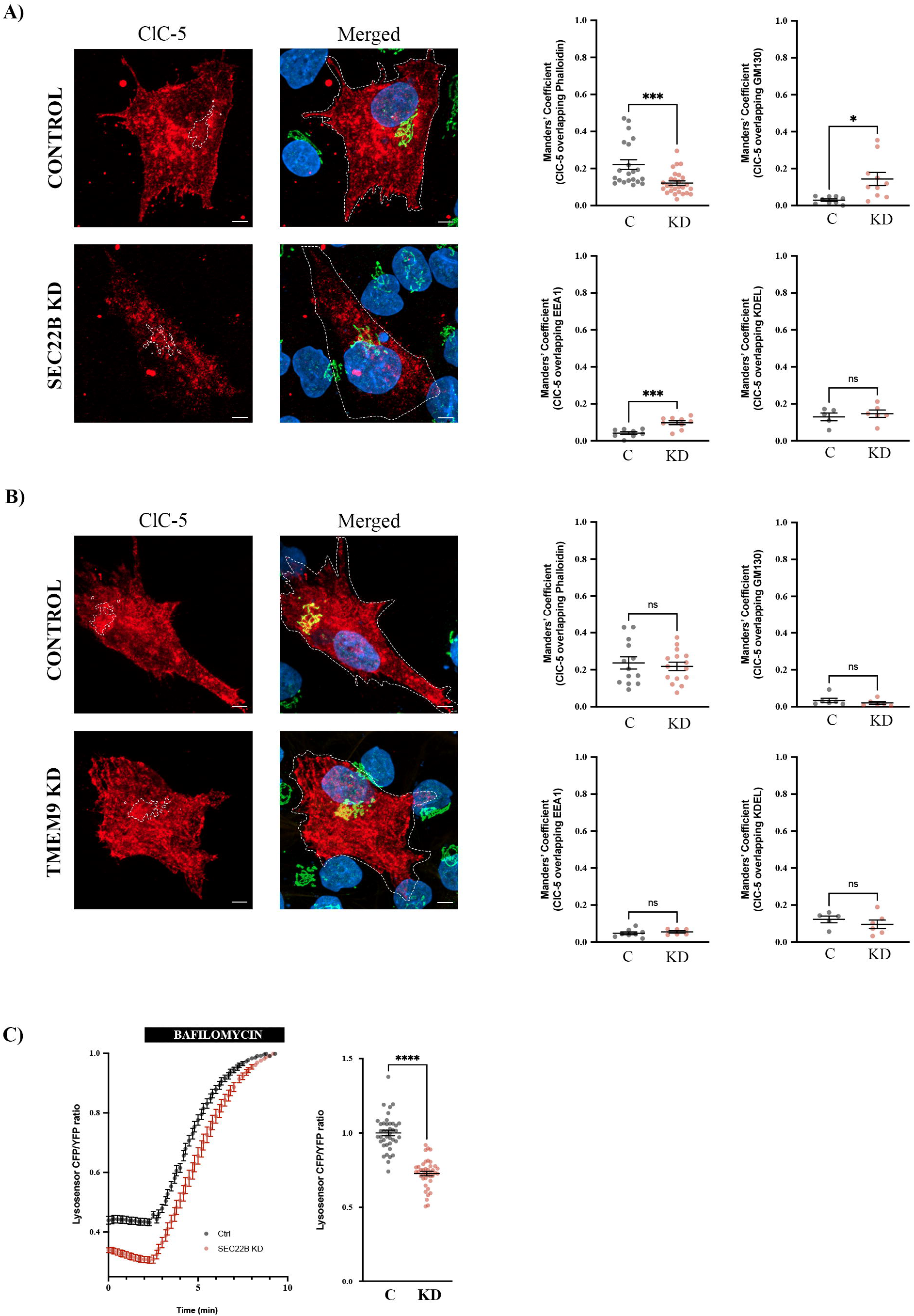
SEC22B deletion impairs ClC-5 trafficking. **A-B)** Representative z-projection images SEC22B KD **(A)** or TMEM9 KD **(B)** and respective control cells, transiently transfected with ClC-5 WT and stained with anti-HA antibody to detect ClC-5. Scale bars: 5 µm. Quantification of ClC-5 colocalization (Manders’ coefficient) with distinct intracellular markers (Membrane: Phalloidin, Golgi: GM130, endosomes: EEA1, ER: KDEL) is provided. Data were calculated from immunofluorescence images using Fiji software (N ≥ 3). Average values ± SEM are plotted as a scatter plot. The y-axis represents Manders’ coefficient of the fraction of ClC-5 overlapping with Phalloidin, GM130, EEA1 or KDEL signal accordingly. **C)** Time course of the CFP/YFP LysoSensor Yellow/Blue DND-160 dye ratio in ClC-5 control and SEC22B KD cells. After 3 min of basal activity, bafilomycin was added as a normalization control. Quantification of the normalized basal CFP/YFP LysoSensor Yellow/Blue DND-160 dye ratio is shown in the left graph. Each dot represents one cell (N ≥ 3). Abbreviations: C: Control cells, KD: SEC22B or TMEM9 KD cells, n.s.: not statistically significant. *p<0.05, **p<0.01, ***p<0.001, ****p<0.0001.

## Discussion

ClC-5, Cl^-^/H^+^ antiporter 5, is crucial for the proper function of the renal proximal tubule. Mutations in this antiporter lead to Dent’s Disease type 1 (DD1), a rare X-linked genetic disorder characterized by proximal tubule dysfunction that progresses to renal fibrosis and kidney failure in males. However, although ClC-5 mutations are the genetic cause of the disease, there is a high phenotypic variability between patients. In order to understand these differences, we have performed an interactome study, identifying several potential interactors of ClC-5. Amongst them, we have confirmed TMEM9 as a regulatory partner of ClC-5, and that disruption of this interaction contributes to some of the features observed in DD1. Importantly, we have identified for the first time an R-SNARE protein called SEC22B to be required for the correct localization of ClC-5 to the plasma membrane. Deletion of this protein impairs trafficking of ClC-5, leading to cellular characteristics similar to those observed in cells carrying ClC-5 mutations.

### 1. Identification of a new class of ClC-5 mutations

A key finding of our study is the demonstration that the ClC-5 V523Δ mutant is accumulated at the Golgi apparatus. Interestingly, other studies have also shown partial Golgi localization for other ClC-5 mutants, especially those affecting the CBS2 domain^25,26^. Our results, therefore, strongly suggest the existence of a distinct group of ClC-5 mutations that cause protein misslocalization and retention at the Golgi apparatus, which we have tentatively named as Class IV. We hypothesize that this accumulation may lead to defects in the secretory pathway, contributing to the phenotypic variability observed in DD1 patients. In addition, this misslocalization could explain why some DD1 patients may present extrarenal manifestations that are more commonly described in Dent’s Disease type 2 (DD2), a condition which is caused by mutations in the inositol polyphosphate-5-phosphatase OCRL, a Golgi-resident protein. OCRL loss-of-function alters PIP_2_ levels, a second messenger essential for many cellular processes such as vesicular transport, therefore impairing intracellular trafficking^15,27,28^. Thus, ClC-5 retention at the Golgi due to specific mutations could also lead to similar defects in this organelle, impacting in the whole secretory pathway. Interestingly, our data revealed that not only Golgi-retained mutants, but also those causing ClC-5 accumulation in the ER or the endosomes alter the secretory pathway of renal proximal tubule epithelial cells. This raises the question whether modification of endolysosomal acidification can affect the organization of the secretory pathway and, thus, the intracellular trafficking, potentially explaining some of DD1 phenotypic characteristics beyond the well-described endocytosis impairment.

### 2. Novel interactors of ClC-5

The results from our interactome analysis revealed several potential interactors of ClC-5, some of them specific of the WT form that were lost in ClC-5 mutants. Interestingly, a large percentage of these interactors were related to the organization of the secretory pathway, protein intracellular trafficking or posttranslational modifications. For instance proteins related to vesicle mediated transport such as SEC22B, SCFD1, or DDOST; or glycosyltransferases like FUT8, or GALNT1. Furthermore, analysis of the potential hits for ClC-5 mutants showed the appearance of new interactors, such as DENND3 for V523Δ mutant, CD2AP for E527D mutant or TCTN2 for the I524K mutant. These distinct novel interactions could help understanding the variability in patients’ progression and DD1 phenotype. Importantly, there were several interactions that were consistently maintained between WT and all ClC-5 mutant forms, including the recently identified as ClC-5 regulatory partner TMEM9.

### 3. TMEM9 modulates ClC-5 function

The proton-transporting V-type ATPase complex assembly regulator TMEM9 has been also shown to be an inhibitory partner for ClC antiporters. Importantly, our results are aligned with those from Planells-Cases et al that revealed the inhibitory role of TMEM9 on ClC antiporters^22^. In this sense, we have confirmed that TMEM9 depletion causes an increase of endolysosomal pH, correlating with the expected effect of removing an inhibitory subunit for ClC-5 function. However, our findings reveal a more complex relationship between ClC-5 and TMEM9, since deletion of the later phenocopies ClC-5 loss-of-function, such as endocytosis defects and epithelial dedifferentiation. We suggest that fine tuning of ClC-5 – TMEM9 levels, interaction and localization is required for the correct acidification of the secretory pathway and, thus, the function of the renal proximal tubule. In fact, strict control of intraorganellar pH is essential for proper organization of the secretory pathway, being around 7 at the ER, decreasing to 6 at the Trans-Golgi to reach 5.2 in secretory vesicles or 4.5 in lysosomes^29,30^. Proper acidification is crucial for protein folding, enzyme activity and also for protein transport and sorting, and glycosylation. Therefore, alterations in the control of acidification, caused by ClC-5 or TMEM9 loss of function, through the secretory pathway can have an impact in cellular processes leading to various pathological conditions.

### 4. SEC22B, a new regulator of ClC-5 trafficking

Notably, our findings showed that both TMEM9 or ClC-5 loss-of-function caused fragmentation of the Golgi apparatus, supporting the idea that fine-regulation of intralumenal acidification is needed for proper organization and function of the secretory pathway. This result permitted to identify the vesicle-trafficking protein SEC22B, a potential interactor of ClC-5, as a possible regulator of their trafficking. SEC22B has been shown by other authors as a R-SNARE required for export and recycling of proteins^23,31^. Our results confirmed that depletion of this R-SNARE caused an alteration of the secretory pathway also in proximal tubule epithelial cells. Notably, SEC22B deletion caused strong retention of ClC-5 at the Golgi apparatus and endosomes, leading to most of the DD1 phenotypic characteristics, such as decreased endocytosis and epithelial markers. Intriguingly, SEC22B depletion also caused higher acidification of endolysosomal pH, similar to the phenotype observed in TMEM9 lacking cells. Furthermore, our findings revealed that SEC22B deletion also impacted in TMEM9 trafficking, reducing its localization at the lysosomes. Therefore, a reasonable explanation is that the higher retention of ClC-5 at the endosomes together with the decreased localization of TMEM9 at the lysosomes caused by SEC22B loss leads to higher acidification of the endolysosomal system. Therefore, our findings suggest that SEC22B is required for the intracellular transport of both ClC-5 and TMEM9 to their final destination, and its aberrant accumulation in other cellular compartments leads to DD1-like characteristics, similar to the phenotype showed by ClC-5 loss-of-function.

In conclusion, we have identified novel interactors of ClC-5, various of them being altered by loss-of-function pathogenic mutations in this antiporter, which could contribute to the development and variability of Dent’s Disease type 1 (DD1) pathophysiology. Amongst them, we propose TMEM9 and SEC22B not only as phenotype modifying genes, but also as potential causative genes for the 30% of patients diagnosed with DD without genetic diagnosis. Consequently, studying mutations or alterations in TMEM9 and SEC22B levels in all DD patients is of high importance. Our results are, therefore, highly relevant for understanding renal tubule function and secretory pathway regulation, as well as clinically for explaining phenotypic variability and providing a potential diagnosis for those DD cases lacking the genetic confirmation.

## Materials and Methods

### Reagents and antibodies

All chemicals were obtained from Sigma-Aldrich (St. Louis, MO) except Alexa Fluor 488 conjugated albumin (Thermo Fisher Scientific, Waltham, MA, USA), LysoSensor Yellow/Blue DND-160 (Thermo Fisher Scientific, Waltham, MA, USA), collagen I, collagen IV and β-catenin antibodies (Abcam, Cambridge, UK), ClC-5 antibody (Genetex, Irvine, CA, USA), SEC22B antibody (Synaptic systems, Göttingen, Germany), and E-cadherin antibody (BD Transduction Labs, Franklin Lakes, NJ, USA). Secondary antibodies for immunofluorescence microscopy and WB were from Life Technologies (ThermoFisher Scientific, Waltham, MA, USA).

### Cell culture and mycoplasma test

Renal proximal tubule epithelial cells RPTEC/TERT1 were obtained from the American Type Culture Collection (#CRL-4031, ATCC®). RPTEC/TERT1 were grown in DMEM: Nutrient Mixture F-12 (1:1, vol/vol) (#31331093; Thermo Fisher Scientific) supplemented with 20 mM HEPES (#15630–080; Gibco), 60 nM sodium selenite (#S9133; Sigma-Aldrich), 5 μg/ml transferrin (#T1428; Sigma-Aldrich), 50 nM dexamethasone (#D8893; Sigma-Aldrich), 100 U/ml penicillin and 100 μg/ml streptomycin (#15240–062; Gibco), 2% FBS (#10270; Gibco), 5 μg/ml insulin (#I9278; Sigma-Aldrich), 10 ng/ml EGF (#E4127; Sigma-Aldrich), and 3 nM triiodothyronine (#T5516; Sigma-Aldrich). Cell culture was maintained at 37 ºC in a humidified atmosphere with 5% CO_2_, and cells were cultured for 10 days to allow for cell differentiation unless otherwise indicated. RPTEC/TERT1 control cells, *CLCN5* knockdown cells (*CLCN5* KD), and knockdown cells overexpressing the ClC-5 WT form (ClC-5 rWT) or the different mutant forms (ClC-5 rV5223del, ClC-5 rE527D, and ClC-5 rI524K) were obtained by stable transduction as described in ^10^. TMEM9 KD, SEC22B KD and respective controls generation is described below. Cells were regularly tested for mycoplasma contamination by PCR as described in ^11^.

### Generation of stable cell lines (shRNA)

For *SEC22B* and *TMEM9* silencing, the MISSION® TRC shRNA transfer vector containing the *SEC22B* shRNA target sequence GAAGTGTTACAACGAGGAGAA (#TRCN0000162016, Sigma-Aldrich) or *TMEM9* shRNA target sequence TGTTGTCTTCTTGGGTCTTTG (#TRCN0000431670, Sigma-Aldrich) were cotransfected with the third-generation packaging vectors VSVG, RTR2 and PKGPIR, which provide the envelope, packaging and reverse-expressing proteins, into HEK293T cells. Viral supernatant was collected, filtered and supplemented with 10% FBS (#10270, Gibco), 1% non-essential amino acids and 8 μg/ml Polybrene Infection / Transfection Reagent (#TR-1003, Sigma-Aldrich) and added to RPTEC/TERT1 cells, after which selection was done using 2 μg/ml puromycin (#ant-pr, InvivoGen).

### Transient transfection HEK293T

HEK293T cells were seeded on 6-well plates at a concentration of 0.2×10^6^ cells/ml. After 24 h, cells were transfected with 0.25 μg of ClC-5 WT plasmid (generated by our group^10^) and 0.25 μg of SEC22B or TMEM9 containing vectors (RC200569 and RC200100, respectively, Origene Global, Rockville, USA) using lipofectamine 3000 (#L3000001, Thermo Fisher Scientific). Lipofectamine Transfection Reagent was prepared using Opti-MEM™ I Reduced Serum Medium, GlutaMAX™ Supplement (#51985091, Gibco, Thermo Fisher Scientific) and Lipofectamine 3000 Reagent (#L3000001, Thermo Fisher Scientific) and incubated for 15 min. After 24 h, cell media was replaced with fresh supplemented DMEM/F-12, GlutaMAX^TM^ medium.

### Transient transfection RPTEC/TERT1

RPTEC/TERT1 cells were cultured in iBIDI plates for 7 days at a concentration of 5×10^4^ cells/ml to ensure cell differentiation at collection time. Following this, SEC22B, TMEM9 or ClC-5 rWT plasmids (described above) were transiently transfected using Lipofectamine^TM^ 3000 (#L3000001, ThermoFisher Scientific) according to manufacturer’s instructions. Briefly, 0.53 µg DNA were added to an Opti-MEM™ I Reduced Serum Medium, GlutaMAX™ Supplement (#51985091, Gibco, Thermo Fisher Scientific) - Lipofectamine^TM^ 3000 dilution, and DNA-lipid complexes were allowed to form for 15 minutes at RT. DNA-lipid complexes were then added to 70-90% confluent cells with FBS and antibiotic-free medium, and were incubated under normal conditions (37 ºC in a humidified atmosphere, 5% CO_2_) for 2 days before being fixed, quenched, and blocked (see Confocal microscopy and immunofluorescence analysis for details). In the case of co-transfected plasmids, 0.25 µg DNA of each were used.

### qRT-PCR analysis (RNA extraction, RT-PCR)

Total RNA was isolated from cells using TRIzol® Reagent (#15596–026, Life Technologies) following the manufacturer’s protocol, and 1000 ng of RNA in a final volume of 20 μl was retro-transcribed using the High-Capacity cDNA Reverse Transcription Kit (#4387406, Applied Biosystems). For each probe to analyse, 20 ng of cDNA was amplified in a 10 μl final volume of reaction mix containing SYBR Green MasterMix (#A25742, Applied Biosystems) or TaqMan MasterMix (#4369016, Applied Biosystems), according to the manufacturer’s instructions. Endogenous levels of CLCN5 mRNA were measured using specific probes targeting the intact shRNA target sequence (Endogenous *CLCN5* primers: 5’-GGGATAGGCACCGAGAGAT-3’ and 5’-GGTTAAACCAGAATCCCCCTGT-3’). Total levels of SEC22B, TMEM9, EFNB1, BMP4, collagen 1 and 4 (*COL1A1* and *COL4A1*) were measured using specific primers (*SEC22B* primers: 5’-TTGCTAACAATGATCGCCCGAG-3’ and 5’-CCCCCTGCTCAATAATGTAGTGAAA-3’; *TMEM9* primers: 5’-CCAGCTGAAGCCAACAAGAGTT-3’ and 5’-GCAGTAGGCCTCCACGTCAT-3’; *EFNB1* primers: 5’-GTATCCTGGAGCTCCCTCAACC-3’ and 5’-GCTTGTAGTACTCATAGGGCC-3’; *BMP4* primers: 5’-CTACTGCAGGGACCTATGGAGC-3’ and 5’-ACGACCATCAGCATTCGGTTAC-3’; *COL1A1* primers: 5’-GTGGTCAGGCTGGTGTGATG-3’ and 5’-CAGGGAGACCCTGGAATCCG-3’; and *COL4A1* primers: 5’-CAGAGATGGTGTTGCAGGAGTG-3’ and 5’-CCTTTGAGCCGCAAGTCGAAAT-3’). Data were normalized against *TBP or GAPDH* using the following primers: *TBP*: 5’-CGGCTGTTTAACTTCGCTTC-3’ and 5’-CAGACGCCAAGAAACAGTGA-3’; *GAPDH*: 5’-GACCCCTTCATTGACCTCAA-3’ and 5’-TTGACGGTGCCATGGAATT-3’.

### Western Blot

Media of RPTEC/TERT1 cells was replaced 24h before collection with fresh medium, after which they were collected and centrifuged at 1,500xg to remove cell debris. Supernatants were then denatured at 95ºC for 5 min with Laemmli buffer. RPTEC/TERT1 cells were washed with 1x PBS and lysed with 1x SET buffer (10 mM Tris-HCl, pH 7.4, 150 mM NaCl, 1 mM EDTA, and 1% SDS). The protein concentration was quantified using the BCA assay (#23225, Thermo Fisher Scientific), and equal amounts of whole cell extracts (20 μg of protein) were resolved in 10% SDS-PAGE gels and transferred to PVDF membranes (#ISEQ00010, Millipore) at 100V for 1 h except when measuring collagen I or IV and fibronectin 1, that they were transferred for 3 h. Membranes were blocked in 5% non-fat dry milk diluted in PBS-T (1x PBS, 0.1% Tween-20) for 1h, and incubated overnight at 4ºC with: HA (1:1000 dilution, #11867423001, Roche), β-tubulin (1:5000 dilution, #T4026, Sigma-Aldrich), collagen I (1:500 dilution, #ab34710; Abcam), collagen IV (1:500 dilution, #ab6586; Abcam), E-cadherin (1:1000 dilution, #610181; BD Transduction Labs), ClC-5 (1:1000 dilution, #GTX53963; Genetex), and fibronectin 1 (1:5000 dilution, #ab45688; Abcam), or blocked in 5% BSA diluted in PBS-T (1x PBS, 0.1% Tween-20) for 1h, and then incubated overnight at 4ºC with SEC22B (1:1000, #186003, Synaptic Systems), MUC1 (1:1000 dilution, #MA1-06503; Thermo Fisher Scientific), cathepsin D (1:1000 dilution, #EPR3057Y; Abcam), HA (1:1000 dilution, #11867423001, Roche), and LAMP-1 (1:1000 dilution, #53-1079-42, Cell Signalling Technology). Membranes were then incubated with the corresponding secondary antibodies (rabbit anti-mouse IgG/HRP, #P0260, Dako, goat anti-rat IgG/HRP, #A9037, Sigma-Aldrich, goat anti-rabbit IgG/HRP, # P0448, Dako, goat anti-Mouse IgG (H+L) 680, #A21109, Thermo Fisher Scientific, goat anti-Rabbit IgG (H+L) 800, #A32730, Thermo Fisher Scientific) at a 1:5000 dilution. Band intensities were visualized with or without chemiluminescence reagent (#WBLUF0500, Millipore) using Odyssey Fc Imaging System (LI-COR), and analysed using FIJI.

### Adhesion assay

RPTEC/TERT1 cells were cultured for 10 days to allow for cell differentiation, and then trypsinized and counted. Forty thousand cells/well were seeded onto two duplicated 96-well plates, with one lane being left without cells for normalization, and allowed to adhere for 1h at 37ºC. Following the incubation period, one of the plates was washed twice with 1x PBS to remove unattached cells, and then cell recovery was allowed for an additional 1h at 37ºC. After this, the tetrazolium salt (XTT) assay (#11465015001, Sigma-Aldrich) was used to measure the amount of cells according to the manufacturer’s instructions. The XTT reagent was added and left to incubate in a dark environment for 4h at 37ºC, after which the absorbance was measured at 490 and 630 nm using the Spark® Tecan microplate reader. Results were expressed as the ratio of XTT values between the washed and unwashed plate, and normalized against the background fluorescence

### Confocal microscopy and immunofluorescence analysis

RPTEC/TERT1 cells were cultured on iBIDIs during 10 days to allow for cell differentiation. The day of the experiment, cells were fixed with 4% PFA for 15 min at RT. Aldehyde groups were quenched in 50 mM 7.4 pH NH_4_Cl/PBS for 30 min, and non-specific binding sites were blocked with 5% BSA in PBS for 60 min. Primary antibodies were diluted in blocking reagent and incubated overnight at 4°C (collagen I [#ab34710; Abcam], collagen IV [#ab6586; Abcam], MUC1 [MA1-06503; Thermo Fisher Scientific], LAMP-1 [#53-1079-72; Invitrogen], KDEL [#Ab10C3; Abcam], β-catenin [#ab32572; Abcam], GM130 [#ab52649; Abcam], vti1a [#V85620; Transduction Technologies], HA [#11867423001; Sigma-Aldrich], p230 [#G88620; Transduction Technologies], EEA1[#3288S; Cell Signalling Technologies], FLAG [#MA191878; Invitrogen], SEC22B [#186003; Synaptic Systems]). Secondary antibodies conjugated with 488 (#A28175; Invitrogen), 568 (#A11011; Invitrogen), and 647 (#A21472; Invitrogen) were diluted in a blocking solution and incubated for 1 h at RT. Cells were incubated with Hoechst 33342 (1:2,000 dilution, #H1399; Invitrogen) or phalloidin (1:200 dilution, #R415; Invitrogen) for 1 h at RT. Fluorescence was visualized in a confocal spectral Zeiss LSM 980 microscope. Images were processing using FIJI software. Manders’ overlap coefficient was calculated using the ImageJ plugin JACoP in a minimum of three different fields (>5 cells/ field) for three independent experiments per condition. JACoP plugin measures colocalization using different indicators (Pearson’s coefficient, overlap coefficient, or Manders’ coefficients) on two images for all z-stacks. To determine the number, and volume of LAMP1-, and EEA1 positive elements, we used the 3D objects counter v2.0 tool from FIJI. All images for each experiment were taken on the same day under the same conditions and the same z-step (0.5 μm), and a minimum of three fields for each experiment (N > 3) were analysed. The threshold was automatically set for each image. DAPI was used to count the number of nuclei per field.

### Endocytosis assay

RPTEC/TERT1 cells were seeded on IBIDI plates and grown for 10 days. To measure albumin uptake, cells were washed with an isotonic buffer (2.5 mM KCl, 140 mM NaCl, 1.2 mM CaCl_2_, 0.5 mM MgCl_2_, 5 mM glucose, and 10 mM HEPES (305 mosmol/litre, pH 7.4 adjusted with Tris)), and incubated with a solution 50 μg/mL Alexa Fluor 488-conjugated albumin in isotonic buffer (#A13100, Thermo Fisher Scientific) for 30 minutes at 37ºC. After the treatment, cells were washed twice with 1x PBS and fixed in 4% PFA for 15 min at RT. Aldehyde groups were quenched in 7.4 pH 50 mM NH_4_Cl/PBS for 30 min, and cells were incubated with Hoechst 33342 (1:2,000 dilution, #H1399; Invitrogen) and phalloidin (1:200 dilution, #R415; Invitrogen) for 1 h at RT to allow visualization of cellular perimeter and cell nuclei. Images were acquired with a 63X objective on a confocal laser scanning microscope (Zeiss LSM 980) and processed using FIJI software.

### Endolysosomal pH determination

Endosomal pH was determined using the LysoSensor Yellow/Blue DND-160 dye (L7545; Thermo Fisher Scientific). RPTEC/TERT1 cells were cultured in IBIDI chambers for 10 days. Cells were washed three times with isotonic solution containing 2.5 mM KCl, 140 mM NaCl, 1.2 mM CaCl_2_, 0.5 mM MgCl_2_, 5 mM glucose, and 10 mM HEPES (305 mosmol/litre, pH 7.4 adjusted with Tris), and then incubated with LysoSensor for 10 min at 37°C. After the incubation period, cells were washed again with isotonic solution and imaged using Leica Thunder Imager 3D Cell Culture Microscope for 3 min without stimulation. Then, cells were treated with vehicle (DMSO) or 100 nM bafilomycin A1 to get a normalization point. Images were acquired with a 5X objective on Leica Thunder Imager 3D Cell Culture Microscope and analysed using FIJI software.

### Sample preparation and TMT labelling

Reduction of disulphide bridges in cysteine containing proteins was performed with dithiothreitol (56°C, 30 min, 10 mM in 50 mM HEPES, pH 8.5). Reduced cysteines were alkylated with 2-chloroacetamide (room temperature, in the dark, 30 min, 20 mM in 50 mM HEPES, pH 8.5). Samples were prepared using the SP3 protocol^32,33^ and trypsin (sequencing grade, Promega) was added in an enzyme to protein ratio 1:50 for overnight digestion at 37°C. Peptides were labelled with TMT6plex Isobaric Label Reagent (ThermoFisher) according the manufacturer’s instructions. Followed by quenching the reaction with 5% hydroxylamine for 15min. RT. Samples were combined for the TMT5plex and for further sample clean up an OASIS® HLB µElution Plate (Waters) was used. Offline high pH reverse phase fractionation was carried out on an Agilent 1200 Infinity high-performance liquid chromatography system, equipped with a Gemini C18 column (3 μm, 110 Å, 100 x 1.0 mm, Phenomenex)^34^.

### Mass spectrometry data acquisition

An UltiMate 3000 RSLC nano LC system (Dionex) fitted with a trapping cartridge (µ-Precolumn C18 PepMap 100, 5µm, 300 µm i.d. x 5 mm, 100 Å) and an analytical column (nanoEase™ M/Z HSS T3 column 75 µm x 250 mm C18, 1.8 µm, 100 Å, Waters). Trapping was carried out with a constant flow of solvent A (0.1% formic acid in water) at 30 µL/min onto the trapping column for 6 minutes. Subsequently, peptides were eluted via the analytical column with a constant flow of 0.3 µL/min with increasing percentage of solvent B (0.1% formic acid in acetonitrile) from 2% to 4% in 4 min, from 4% to 8% in 2 min, then 8% to 28% for a further 37 min, and finally from 28% to 40% in another 9 min. The outlet of the analytical column was coupled directly to a QExactive plus (Thermo) mass spectrometer using the proxeon nanoflow source in positive ion mode.

The peptides were introduced into the QExactive plus via a Pico-Tip Emitter 360 µm OD x 20 µm ID; 10 µm tip (New Objective) and an applied spray voltage of 2.1 kV. The capillary temperature was set at 275°C. Full mass scan was acquired with mass range 375-1200 m/z in profile mode with resolution of 70000. The filling time was set at maximum of 10 ms with a limitation of 3×10^6^ ions. Data dependent acquisition (DDA) was performed with the resolution of the Orbitrap set to 17500, with a fill time of 50 ms and a limitation of 2×10^5^ ions. A normalized collision energy of 32 was applied. Dynamic exclusion time of 30 s was used. The peptide match algorithm was set to ‘preferred’ and charge exclusion ‘unassigned’, charge states 1, 5 - 8 were excluded. MS2 data was acquired in profile mode. The mass spectrometry proteomics data have been deposited to the ProteomeXchange Consortium via the PRIDE^35^ partner repository with the dataset identifier PXD070231.

### Data analysis

The protein.txt output files of IsobarQuant^36^ were processed with the R programming language (ISBN 3-900051-07-0). Only proteins that were quantified with at least two unique peptides in at least two out of three replicates were considered for the analysis. The raw summed tmt reporter ion signals (‘signal_sum’ columns) were used and first cleaned for potential batch-effects using the ‘removeBatchEffect’ function of the limma package^37^. The resulting data were normalized with a variance stabilization normalization (vsn)^38^. Missing values were imputed with the ‘impute’ function (method = ‘knn’) of the Msnbase package^39^. Limma was employed again to test for differential expression between different conditions. The replicate factor was included in the linear model of limma. Proteins were considered ‘hit’ with a false discovery rate (fdr) smaller 5 % and a fold-change greater 100 % and considered ‘candidate’ with a false discovery rate smaller 20 % and a fold-change greater 50 %.

### Statistical analysis

Continuous data are presented as means ± SEM, and outliers were removed using the Grubb’s test (alpha□= □0.05). In all cases, a D’Agostino–Pearson omnibus normality test was performed before any hypothesis contrast test. Statistical analysis and graphics were performed using GraphPad Prism 9 (RRID:SCR_002798) software. For data that followed normal distributions, we applied either a t test or one-way ANOVA followed by Tukey’s post hoc test. For data that did not fit a normal distribution, we used Mann– Whitney’s unpaired t test and non-parametric ANOVA (Kruskal–Wallis) followed by Dunn’s post hoc test. Criteria for a significant statistical difference were as follows: *p < 0.05, **p < 0.01, ***p < 0.001, ****p < 0.0001.

## Acknowledgments

We thank the patient advocacy group ASDENT (Asociación de la Enfermedad de Dent, www.asdent.es) for continuously supporting our group, and also the Dent Disease Foundation (DD Foundation, www.dentdisease.org) for promoting international cooperation in Dent Disease Research. We also thank all members of the Renal Physiopathology Group for valuable discussions. Fluorescence microscopy was performed at the High Technology Unit (HTU) at the Vall d’Hebron Research Institute (VHIR). This work reflects only the authors’ views, and the EU Community is not liable for any use that may be made of the information contained therein. This work was supported in part by Dent’s Disease Patients Association ASDENT and grants from Ministerio de Ciencia e Innovación (SAF201459945-R and SAF201789989-R to A Meseguer), Instituto de Salud Carlos III (PI22/00741 to G. Cantero-Recasens, co-financed by the European Union), the SENEFRO foundation (SEN2019 to A. Meseguer), the Red de Investigación Renal REDinREN (RD16/0009/0030), the RICORS20240 (RD24/0004/ 0031 financed by Instituto de Salud Carlos III and co-financed by the European Union), the Catalan Agency AGAUR (2021SGR01600), and the Mizutani Foundation (REF230040 to G. Cantero-Recasens). M.D. and E.S. were supported by the generous contribution of ASDENT Patients’ Association. A.C-P. was a recipient of the Ph.D. VHIR fellowship 2024 Call. The Renal Physiopathology research group holds the Quality Mention from the Generalitat de Catalunya.

## Author contributions

M Durán: formal analysis, investigation, methodology, and writing—review and editing.

A Casal-Pardo: formal analysis, investigation, methodology, and writing—review and editing.

E Sarró: conceptualization, investigation, and writing—review and editing

F Stein: formal analysis, and writing—review and editing.

S Cases-Palau: investigation, and writing—review and editing.

C Garcia: investigation, and writing—review and editing.

G Ariceta: investigation and writing—review and editing.

A Meseguer: conceptualization, resources, formal analysis, supervision, funding acquisition, and writing—review and editing.

G Cantero-Recasens: conceptualization, formal analysis, supervision, funding acquisition, validation, investigation, methodology, and writing—original draft, review, and editing.

## Competing interests

The authors declare that they have no conflict of interest.

## Supplementary figures

**Supplementary figure 1**. <u>DD1 cell lines characterization.</u> **A)** Endogenous *CLCN5* mRNA levels normalized to values of *GAPDH* from control (Ctrl), *CLCN5* KD (KD), ClC-5 rWT (rWT), ClC-5 rV523del (rV523Δ), ClC-5 rE527D (rE527D), and ClC-5 rI524K (rI524K) cells. All values are represented as a relative value compared with control cells. Average values ± SEM are plotted as a scatter plot with a bar graph (N ≥ 3). **B)** Cell lysates from ClC-5 rWT (rWT), ClC-5 rV523del (rV523Δ), ClC-5 rE527D (rE527D), and ClC-5 rI524K (rI524K) cells were analysed by Western blot with an anti-HA to test expression levels. Tubulin was used as a loading control. Quantification of ClC-5 protein levels is provided as a scatter plot (N ≥ 3). **C)** Quantification of ClC-5 colocalization (Manders’ coefficient was calculated using FIJI software) with different intracellular markers: KDEL: Endoplasmic Reticulum (ER), vti1a: Golgi Apparatus (GA), p230: Trans-Golgi Network (TGN), EEA1: Endosomes. Average values ± SEM are plotted as a scatter plot. The y-axis represents Manders’ coefficient of the fraction of ClC-5 (HA) overlapping with KDEL, vti1a, p230 and EEA1 signal respectively (N ≥ 3). **D)** Number and volume of endosomes was quantified from immunofluorescence images stained with anti-EEA1 antibody using Fiji software (N ≥ 3). Abbreviations: Ctrl: control; KD: CLCN5 KD; rWT: ClC-5 rWT; rV523del: ClC-5 rV523del; rE527D: ClC-5 rE527D; rI524K: ClC-5 rI524K. *p<0.05, **p<0.01, ***p<0.001, ****p<0.0001.

**Supplementary figure 2**. <u>TMEM9 in ClC-5 mutant cell lines.</u> **A)** Representative z-projection images of overexpressed TMEM9 in ClC-5 rWT (rWT), ClC-5 rV523del (rV523Δ), ClC-5 rE527D (rE527D), and ClC-5 rI524K (rI524K) cells and stained with anti-FLAG antibody (TMEM9), anti-LAMP1 (lysosomes) and Hoechst 33342. **B)** Endogenous *TMEM9* mRNA levels normalized to values of *GAPDH* from control (Ctrl) and TMEM9 KD (KD) cells. All values are represented as a relative value compared with control cells. Average values ± SEM are plotted as a scatter plot (N ≥ 3). Abbreviations: Ctrl: Control, KD: TMEM9 KD. ****p<0.0001.

**Supplementary figure 3**. <u>Activation of β-catenin pathway in TMEM9 KD cells.</u> **A)** Immunofluorescence single planes of control and TMEM9 KD cells stained with anti-β-catenin (green), phalloidin (red), and Hoechst 33342 (blue). The nucleus perimeter is demarcated with dotted white lines to visualize β-catenin nuclear translocation. Scale bars: 5 μm. β-Catenin levels at the nucleus (left graph) or at the membrane (right graph) normalized to total levels were quantified from immunofluorescence images. N>3. **B)** *BMP4*, and *EFNB1* mRNA levels (normalized to *TBP* or *GAPDH* levels) from control (C, black dots), and TMEM9 KD (KD, pink dots) cells. **C-D)** *COL1A1* (Col I) **(C)** and *COL4A1* (Col IV) **(D)** mRNA levels (normalized to *TBP* levels) from control (black dots) or TMEM9 KD (pink dots) cells. Abbreviations: C: control, KD: TMEM9 KD, ns: not statistically significant. *p<0.05, **p<0.01.

**Supplementary figure 4**. <u>Interaction between SEC22B and ClC-5.</u> **A)** SEC22B co-immunoprecipitation with the different forms of ClC-5 (WT, rV523Δ, rE527D and rI524K). **B)** Immunofluorescence z-stack single planes of ClC-5 rWT (rWT), ClC-5 rV523del (rV523Δ), ClC-5 rE527D (rE527D), and ClC-5 rI524K (rI524K) cells stained with anti-FLAG (SEC22B, green), anti-KDEL (ER marker, red), and Hoechst 33342 (Blue). Scale bars: 5 µm. **C)** Endogenous *SEC22B* mRNA levels normalized to values of *TBP* from control (Ctrl) and SEC22B KD (KD) cells. All values are represented as a relative value compared with control cells. Average values ± SEM are plotted as a scatter plot (N ≥ 3). **D)** Cell lysates from control and SEC22B KD cells were analysed by Western blot with an anti-SEC22B to test expression levels. Tubulin was used as a loading control. Quantification of SEC22B protein levels is provided. Abbreviations: Ctrl: control; KD: CLCN5 KD; rWT: ClC-5 rWT; rV523del: ClC-5 rV523del; rE527D: ClC-5 rE527D; rI524K: ClC-5 rI524K. ****p<0.0001.

**Supplementary figure 5**. <u>β-catenin pathway in SEC22B KD cells.</u> **A-B)** *COL1A1* (Col I) **(A)** and *COL4A1* (Col IV) **(B)** mRNA levels (normalized to *TBP* or *GAPDH* levels, respectively) from control (black dots) or SEC22B KD (pink dots) cells. **C)** Immunofluorescence single planes of control and SEC22B KD cells stained with anti-β-catenin (green), phalloidin (red), and Hoechst 33342 (blue). The nucleus perimeter is demarcated with dotted white lines to visualize β-catenin nuclear translocation. Scale bars: 5 μm. β-Catenin levels at the nucleus (top graph) or at the membrane (bottom graph) normalized to total levels were quantified from immunofluorescence images. N ≥ 3. D) *BMP4*, and *EFNB1* mRNA levels (normalized to *TBP* or *GAPDH* levels) from control (C, black dots), and SEC22B KD (KD, pink dots) cells. Abbreviations: C: control; KD: SEC22B KD ns: not statistically significant.

**Supplementary figure 6**. <u>Effect of SEC22B and TMEM9 depletion in ClC-5 levels.</u> **A)** Colocalization between TMEM9 (anti-FLAG) and LAMP1 (lysosomes) for Control (Ctrl) and SEC22B KD (KD) cells transfected with TMEM9 was calculated from immunofluorescence images by Manders’ coefficient using FIJI (N ≥ 3). Average values ± SEM are plotted as a scatter plot with a bar graph. The y-axis represents Manders’ coefficient of the fraction of TMEM9 (anti-FLAG) overlapping with LAMP1. **B)** Endogenous *CLCN5* mRNA levels normalized to values of *TBP* from TMEM9 and SEC22B KD cells and respective controls. All values are represented as a relative value compared with respective control cells. Average values ± SEM are plotted as a scatter plot with a bar graph (N ≥ 3). **C)** Cell lysates from TMEM9 KD, SEC22B KD and control cells were analysed by Western blot with an anti-ClC-5 to test expression levels. Tubulin was used as a loading control. Quantification of ClC-5 protein levels is provided as a scatter plot (N≥15). Abbreviations: Ctrl: control, KD: SEC22B or TMEM9 KD. *p<0.05, **p<0.01.

## Notes

### Competing Interest Statement

The authors have declared no competing interest.

