## Supplementary figures and images for "TMEM9 and SEC22B interact with ClC-5 to shape renal proximal tubule function and Dent’s Disease type I pathogenesis"

### Supplementary Figure 1

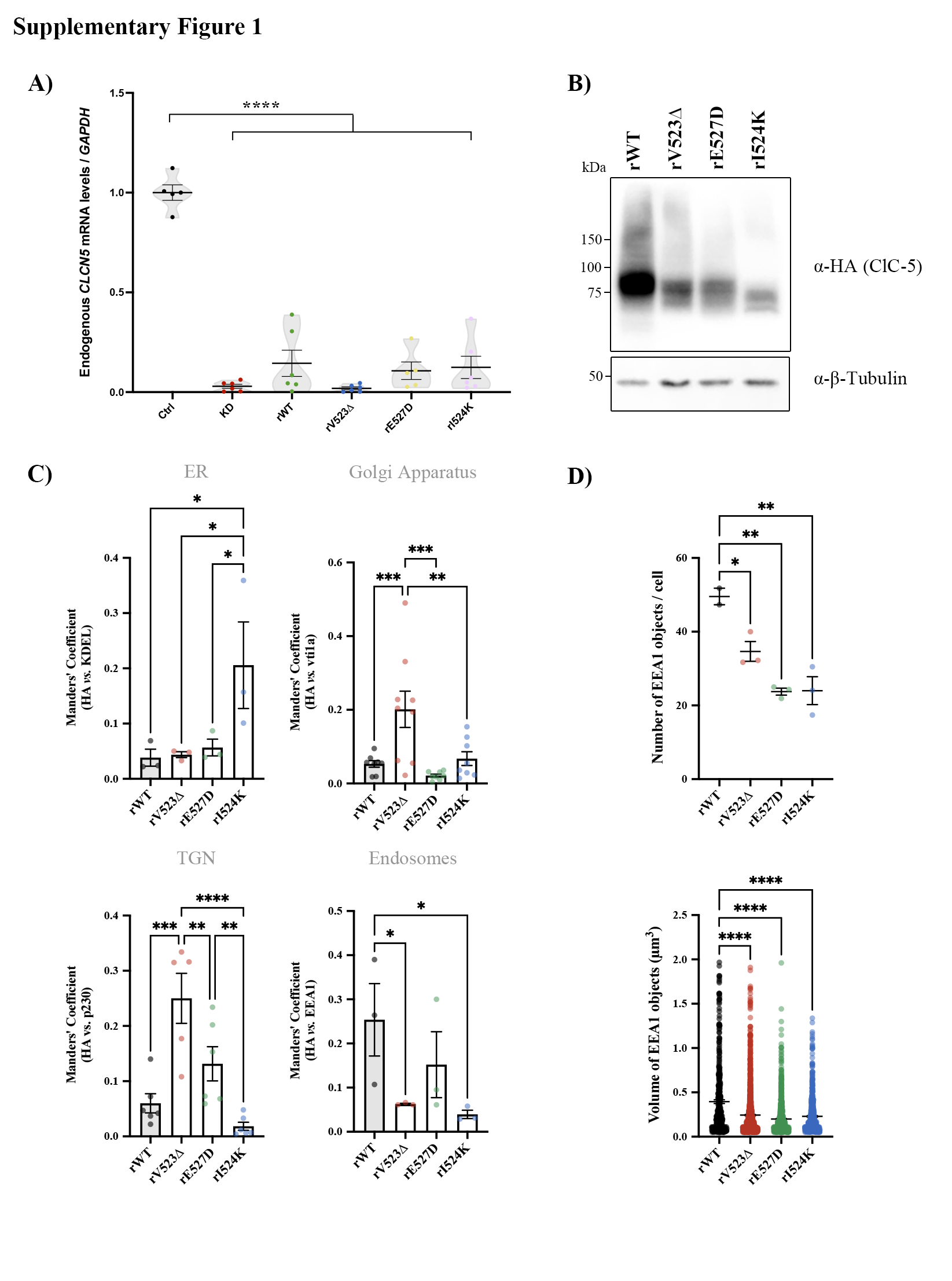

### Supplementary Figure 2

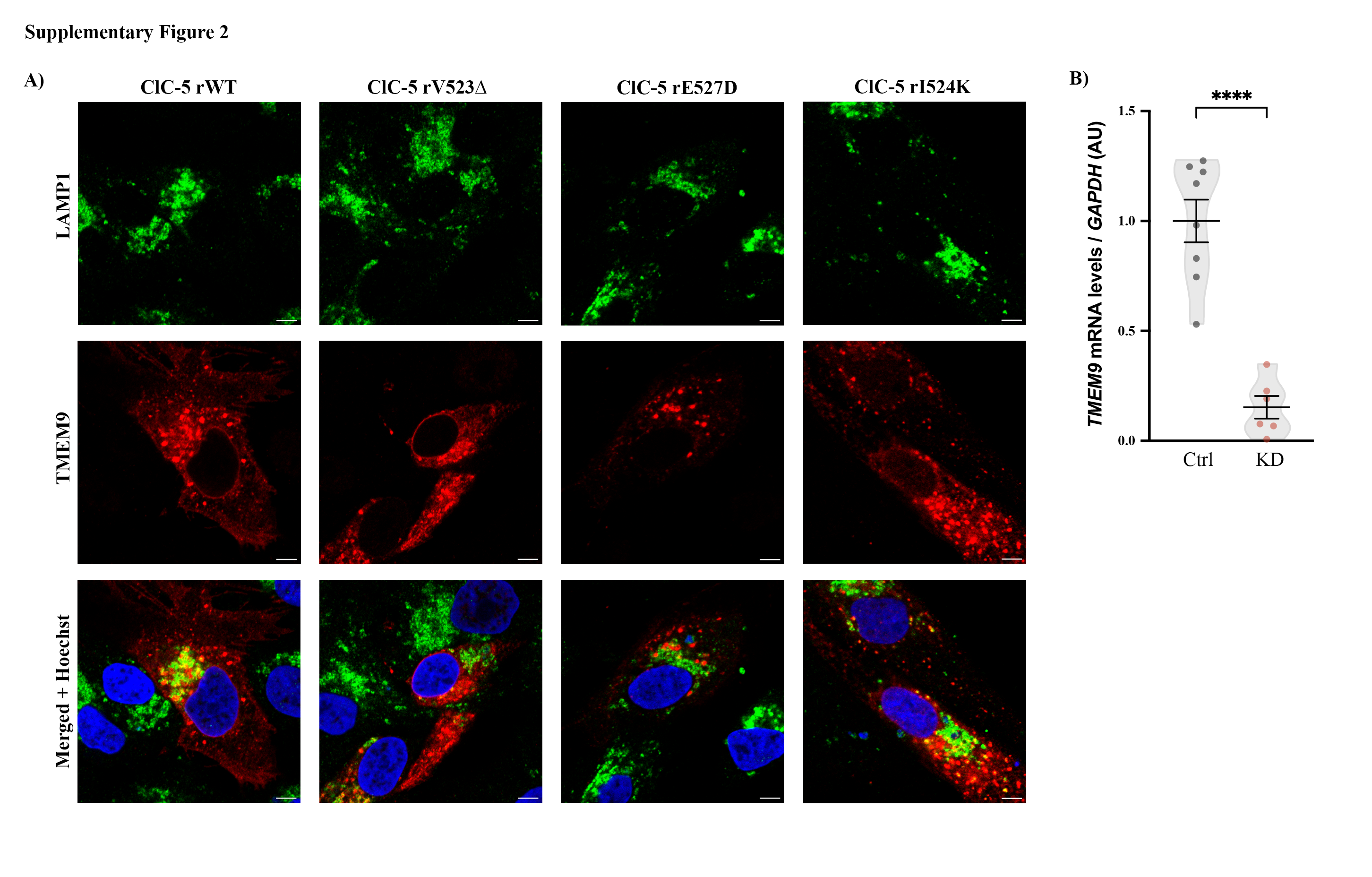

### Supplementary Figure 3

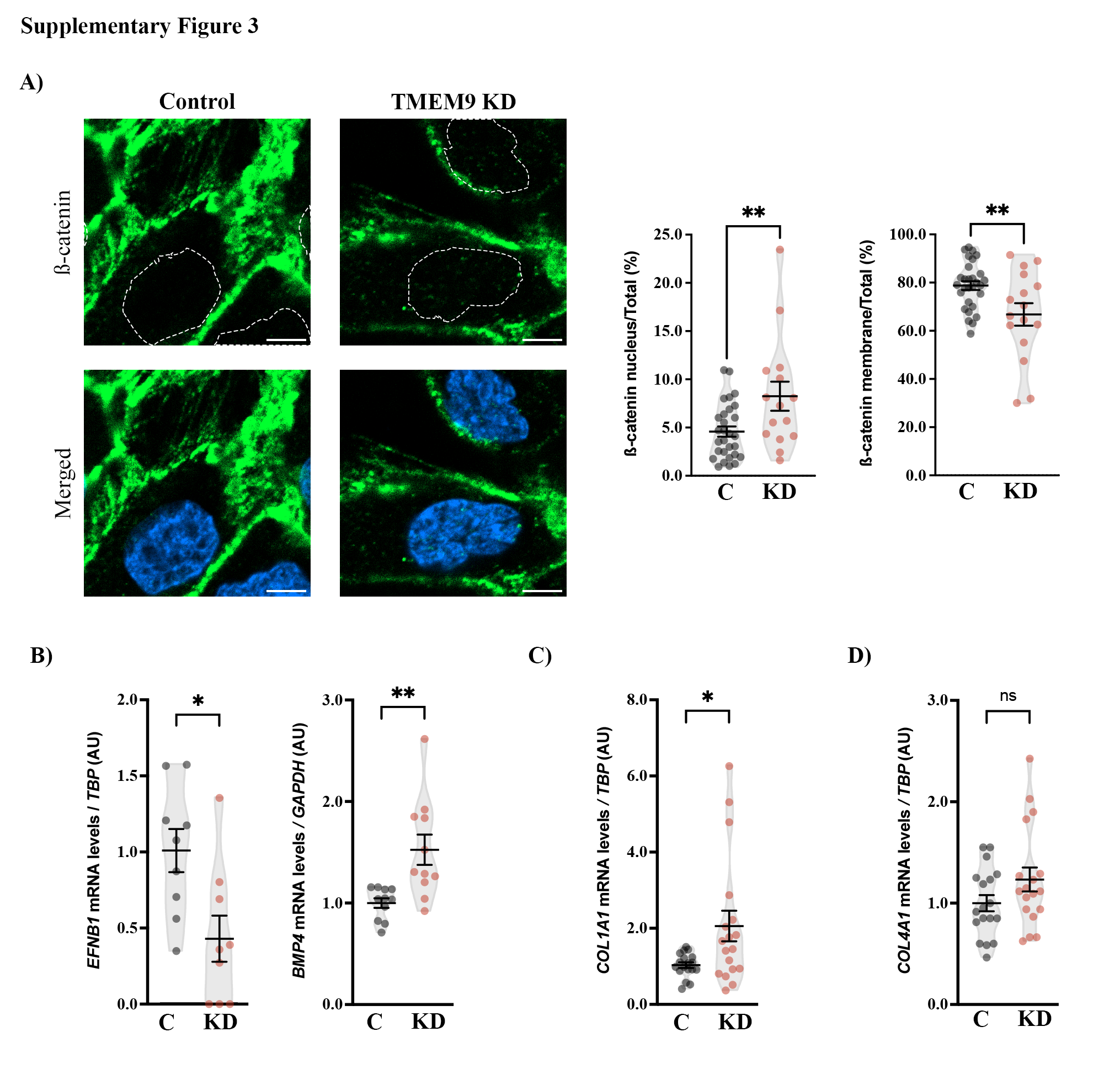

### Supplementary Figure 4

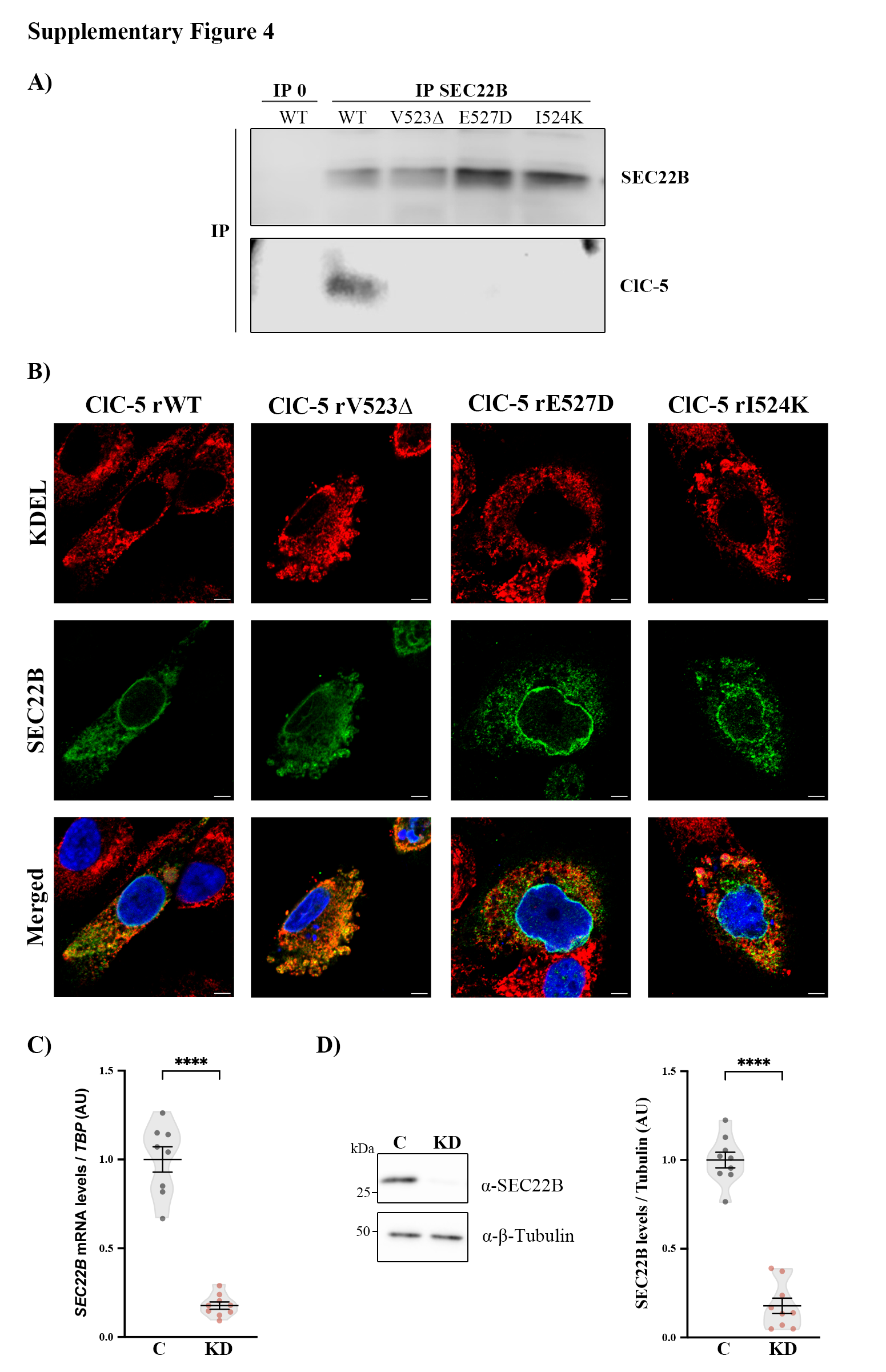

### Supplementary Figure 5

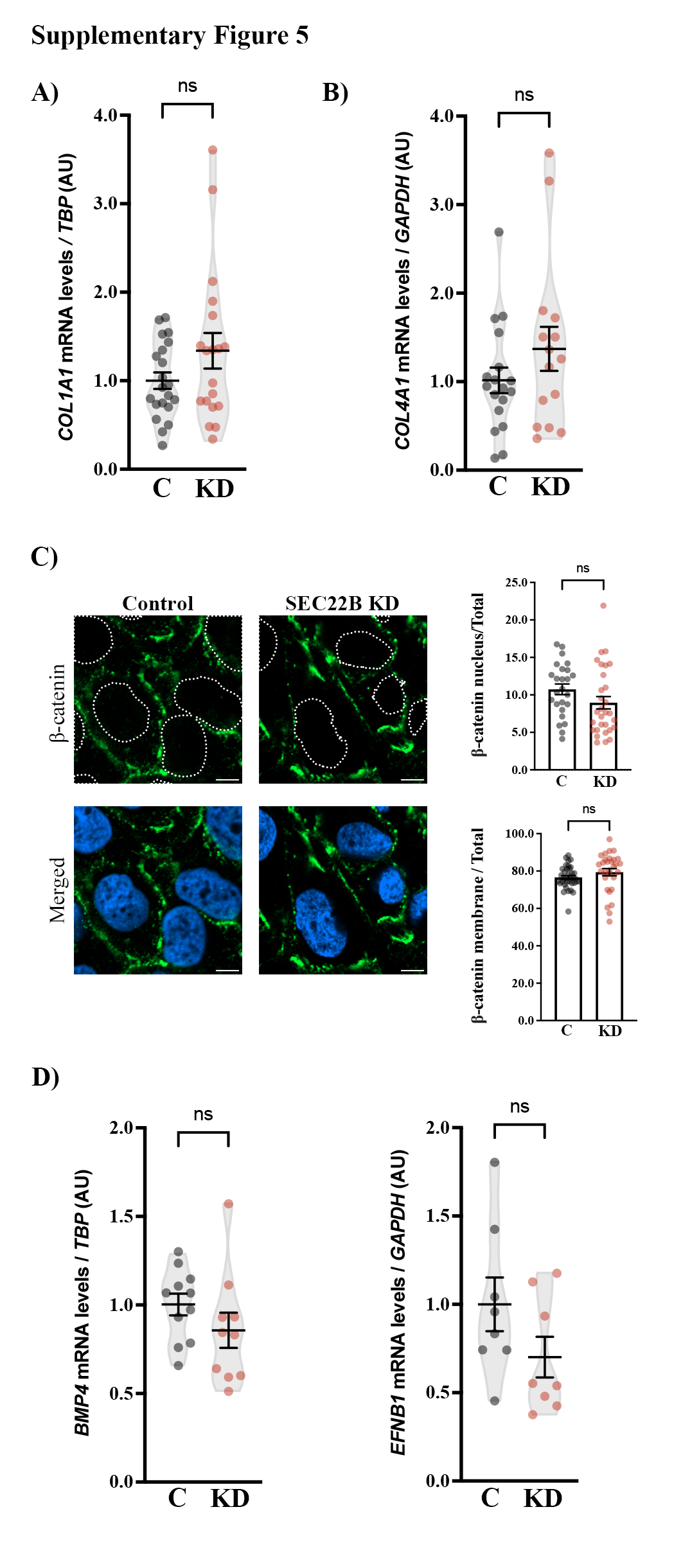

### Supplementary Figure 6

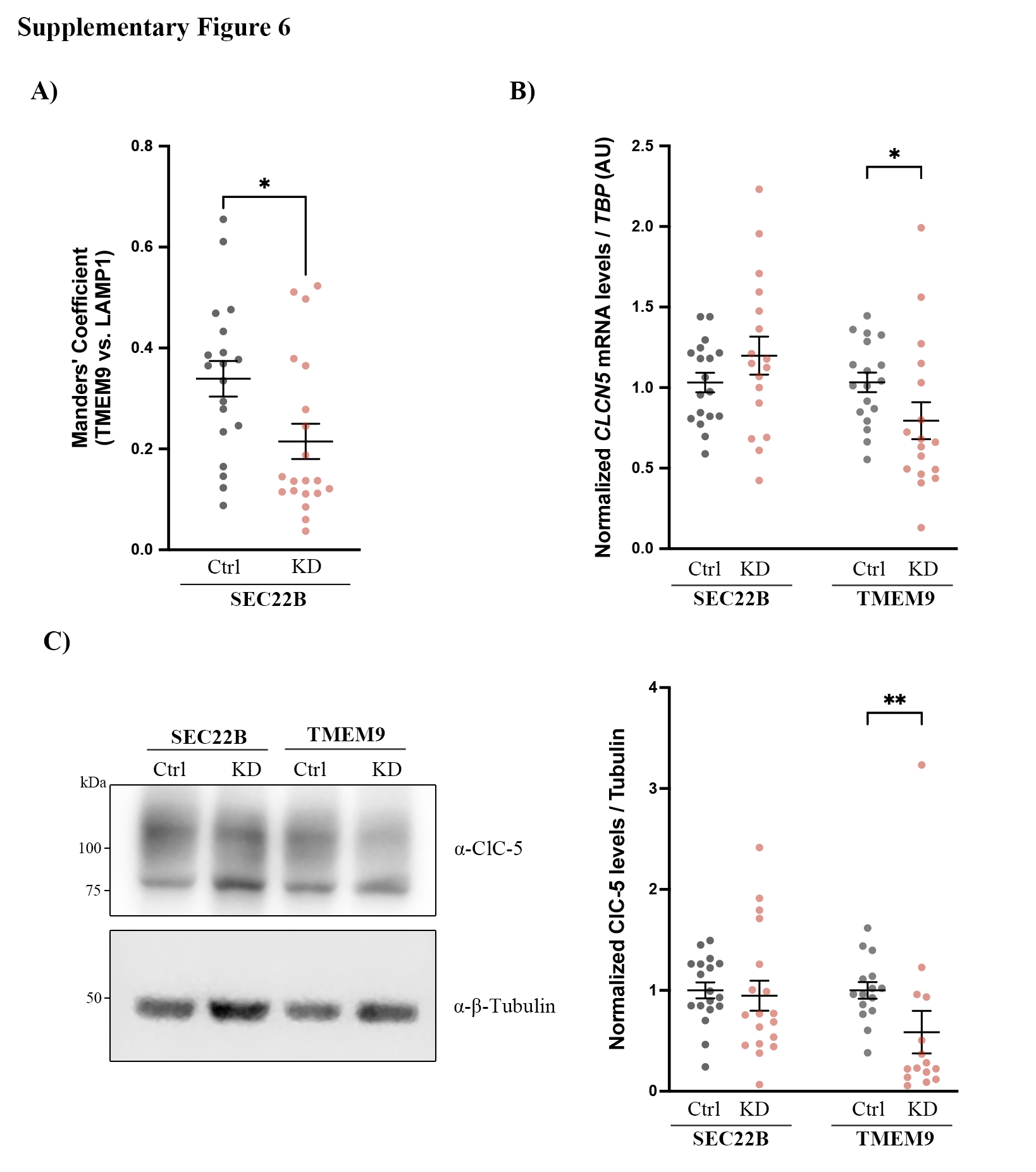
